# Chlorophyll binding to Cytochrome b6f precedes photosystem I and II in barley etioplasts

**DOI:** 10.64898/2026.08.14.744623

**Authors:** Ann Kristin Vatland, Janine Arnold, Dmitry Shevela, Veronika Reisinger, Astrid Mork-Jansson, Bernd Müller, Behzad Heidari, Lutz Andreas Eichacker

**Affiliations:** University of Stavanger, Richard Johnsensgate 4, 4021 Stavanger, Norway; Department of Chemistry, Chemical Biological Centre, Umeå University, S-90187 Umeå, Sweden; Sandoz GmbH, Biochemie-Strasse 10, 6250 Kundl, Austria; Bruker Daltonics GmbH & Co. KG, Fahrenheitstr. 4, 28359 Bremen, Germany

**Keywords:** Chlorophyll *a*, Cytochrome b₆f, light-harvesting-like protein, mass spectrometry, photosystem, plastid, chlorophyll biosynthesis, LIL3 chaperone, etioplast, deetiolation

## Abstract

Chlorophyll (Chl) is essential for oxygenic photosynthesis, binding to membrane proteins for light harvesting and electron transfer. In angiosperms, Chl synthesis is halted in darkness, preventing accumulation of Chl-binding photosynthetic complexes in etioplasts. However, etioplasts assemble a dimeric Cytochrome b6f (Cyt b6f) complex, uniquely binding protochlorophyll (Pchl), the esterified derivative of protochlorophyllide (Pchlide). This indicates an evolutionarily conserved structural or functional role for Pchl distinct from the Chl bound in Cyt b6f in chloroplasts. Here we show that upon light-induced Chl synthesis in-vivo and in-vitro, Chl accumulation in Cyt b6f dimers precedes photosystems I and II. We find that chlorophyllide and Chl bind to the light-harvesting-like protein 3 (LIL3), supporting a role for LIL3 in early Chl allocation that extends its described role in stabilizing geranylgeranyl reductase. We determine a dissociation constant of 246.6 ± 37 nM for Chlide binding to LIL3 in-vitro and show that Cyt b6f monomers and LIL3 co-migrate with Chlide in native PAGE, whereas Cyt b6f dimers and LIL3 co-migrate with Chl. These results indicate that Chlide binding to LIL3 chaperones esterification to Chl and reduction of geranylgeraniol, and that Chl release with Cyt b6f dimerization prioritizes Chl binding to Cyt b6f assembly during de-etiolation.

## Introduction

Oxygenic photosynthesis is an essential metabolic process sustaining heterotrophic life on Earth (Blankenship 2026). The process is built around chlorophyll (Chl), a tetrapyrrole derivative arranged in membrane proteins for light-harvesting and redox chemistry (Croce and Van Amerongen 2014). Chl is bound in three major complexes in the thylakoid membrane of chloroplasts: Photosystem I (PSI) (Nelson and Yocum 2006), Photosystem II (PSII) (Umena et al. 2011), and light-harvesting complexes (LHC) (Wei et al. 2016). These complexes enable efficient energy capture and electron transfer, with PSI and PSII driving linear electron flow for NADPH and ATP production, while LHCs optimize light absorption under varying conditions (Croce and Van Amerongen 2014).

Interestingly, the Cytochrome b6f dimer (Cyt b6f) (Malone et al. 2019), which is key for regulating light-dependent electron and proton flow in photosynthesis (Tikhonov 2014), binds a single chlorophyll per monomer (Dashdorj et al. 2005). This Chl molecule is unique to oxygenic photosynthetic organisms (Ermakova et al. 2024) and is positioned near the Qₚ site, potentially modulating plastoquinone binding and electron flow (Hasan et al. 2014). However, in non-photosynthetic etioplasts, protochlorophyll (Pchl) and not Chl was isolated with the dimeric Cyt b6f complex, indicating that the Cyt b6f complex is already assembled and enzymatically active in etioplasts, and that both chlorins serve a particular structural or functional role in etioplasts and chloroplasts (Reisinger et al. 2008a; Tikhonov 2014; Malone et al. 2019).

During de-etiolation, monomeric PSII complexes assemble with Chl, bind Mn²⁺ and become capable of light-dependent O₂ evolution within about 1 h of illumination, while the Cyt b6f complex is already present in the etioplast (Reisinger et al. 2008a; Shevela et al. 2016). These findings raised the question how Chl binding during Cyt b6f biogenesis is regulated. Biosynthesis of Chl is tightly regulated through feedback inhibition, retrograde signaling, and the light-dependent reduction of protochlorophyllide (Pchlide) to chlorophyllide (Chlide) by light-dependent protochlorophyllide oxidoreductase (LPOR) (Wang et al. 2024). In darkness, ring D of the chlorin structure in Pchlide remains oxidized, conferring lower absorbance and esterification yields by chlorophyll synthase (ChlG) relative to Chlide (Hollingshead et al. 2022). Also, hydrogenation of geranylgeraniol (GG) to phytol (PY) in Pchl is incomplete, yielding higher amounts of partially hydrogenated forms (DHGG, THGG) (Mork-Jansson et al. 2015a).

We previously identified LIL3, a member of the light-harvesting-like (LIL) protein family, as binding Chl within the first seconds of de-etiolation (Reisinger et al. 2008b). LIL3 has since been implicated in Chl biosynthesis by stabilizing geranylgeranyl reductase (GGR), facilitating tocopherol synthesis, and interacting with phytoene synthase for carotenoid biosynthesis (Tanaka et al. 2010; Takahashi et al. 2014; Mork-Jansson et al. 2015a; Hey et al. 2017; Kodru et al. 2024). Cyanobacterial homologs such as HliD bind Chl, associate with chlorophyll synthase (ChlG), and were reported to deliver Chl to photosystems (Chidgey et al. 2014; Proctor et al. 2020). We previously identified amino acids involved in chlorin interaction with LIL3 and proposed how LIL3 dimerization could support a chaperone-like function in phototransformation and esterification (Mork-Jansson et al. 2015a; Mork-Jansson and Eichacker 2018).

Here, we investigated the allocation of Chl to Cyt b6f during the onset of angiosperm deetiolation. We propose that the function of LIL3 may mirror HliD in photosystem biogenesis in cyanobacteria and show in-vitro that Chlide binding to LIL3 results in co-migration with the Cyt b6f monomer, and Chl binding to LIL3 results in co-migration and accumulation of Chl in Cyt b6f dimers. In parallel, we show the Chl dependent assembly of photosystem complexes; however, we find accumulation of Chl in these complexes only after 1 h of deetiolation.

## Materials and Methods

Etioplasts were isolated from 4.5-day-old etiolated barley seedlings (*Hordeum vulgare* L. cultivar Steffi; Saatzucht Ackermann & Co, Irlbach, Germany) grown at 25°C in darkness, as previously described (Eichacker et al. 1996). Membranes were prepared according to (Reisinger et al. 2008a).

### Chl and protein synthesis and supplementation of Chl *a*

Synthesis of etioplast encoded protein was monitored using 35S-Methionine (Eichacker et al. 1996). For synthesis of Chl in the absence of effective protein synthesis, isolated etioplasts (5* 10^7^) were illuminated with white light for 10 s or membranes were supplemented with 0.08 - 1 nmol of Chlide a or Zn-pheide a in the absence or presence of 2.77–5.55 nmol GGPP (Sigma-Aldrich, St Louis, MO, USA) and of 5 mM NADPH (Sigma) for 10 to 80 min at 0 °C or 25 °C. For extraction of membrane protein complexes, reactions were diluted by addition of 1 reaction volume TMK buffer (10 mM Tris-HCl, 10 mM MgCl_2_, 20 mM KCl, pH8.5) and incubated for 10 min on ice, membranes were resuspended and concentrated twice and recovered by centrifugation at 5200xg, for 3 min at 10 °C. The membrane sediment was solubilized for LN-PAGE analysis (see Native PAGE).

For supplementation of etioplast membrane with Chl a, Chl a (from *Anacystis nidulans*; Sigma-Aldrich) was dissolved in 80% acetone/10 mM HEPES, pH 8, and concentrations were determined spectrometrically (Porra et al. 1989). Chl was transferred to the 80% acetone extract of etioplast membranes from which proteins had been removed by precipitation overnight at – 20 °C. The acetone extract was mixed with a corresponding quantity of dissolved Chl and subsequently dried using a speed vacuum dryer and stored in an N_2_ atmosphere. Lysed etioplasts (10^8^ plastids) were used to resuspend the dried extract containing 0.6 to 3 nmol Chl as indicated in the experiments. The incubation time varied from 10 min to 360 min and membranes were recovered by centrifugation at 7000 rpm for 3 min. Chl distribution in the membrane protein complexes was determined after solubilization by LN-PAGE. Alternatively, 10^8^ etioplast membranes were directly supplemented with 1 nmol Chl a in 80% acetone to yield a final concentration of 0.7 nmol and < 3% % acetone (v/v). Etioplast membranes (10^7^) were incubated from 1 to 360 min at RT in the absence or presence of 5 mM NADPH. Chlide was generated by freezing dark-grown seedlings on dry ice for 2 min, followed by 2-s illumination, and pigment extraction in 80% acetone at – 20 °C using a blender. Concentrations were determined using extinction coefficients of ε = 76.79 mM⁻¹ cm⁻¹ at 663 nm (Porra et al. 1989). All work on etiolated samples were conducted in green safelight to minimize photoconversion by POR (Eichacker et al. 1996).

### Native-PAGE and 2D SDS-PAGE

Etioplast membranes (10^8^ plastids) were solubilized in 4.6 mM digitonin (10 °C, 10 min), followed by 5 mM n-dodecyl-β-D-maltoside. For BN-PAGE, Samples were centrifuged (20,100 × g, 10 min) and supplemented with 10% glycerol. Native complexes were separated by LDS-native PAGE (LN-PAGE) over night at constant 35V (Reisinger et al. 2008b; Arnold et al. 2014). Alternatively, etioplasts proteins were radiolabelled using 75 µCi ^35^S-Methionine with a specific activity >1000 Ci/mMol, radiolabel was chased by by 20 mM L-Methionine in the presence of 15 µM Lincomycin and proteins were separated by BN-PAGE (Reisinger and Eichacker 2008). For 2D analysis, LN-PAGE strips were incubated in 150 µl LDS buffer (50mM MES, 50 mM TrisBase, 0.1 % LDS, 1 mM EDTA) for 20 min at RT and separated by LDS-PAGE at constant 200V. Proteins were identified by immunoblotting with anti-Cyt b6(1:10,000; Agrisera AS03-034) and anti-LIL3 (1:7,500; Agrisera, custom against CQSTWQDDSTSGPKK). Secondary HRP-conjugated anti-rabbit IgG (Agrisera) was used for chemiluminescent detection. In-gel heme staining of Cyt b6f complexes was conducted in 6.3 mM TMBZ (3,3’,5,5’-tetramethylbenzidine), 0.25 M sodium acetate buffer, pH 5.0, 30% hydrogen peroxide (H_2_O_2_). Fluorescence was recorded and processed using hardware and software for the Typhoon Trio and Image Quant (GE Healthcare, Buckingham, GB), and the Odyssey CLx (Biotechnology GmbH, Bad Homburg).

### Pigment analysis

Pigments were extracted from etioplast membranes and from protein complexes separated in polyacrylamide gels. A plastid count equivalent basis was used for loading of gels and TLC plates and plastid count numbers of 1-2.5*10^7^ plastids were typically applied. Membranes were extracted in 80% acetone/10 mM Hepes-KOH pH 8 (v/v) at −20 °C, overnight.

Precipitated proteins were sedimented by centrifugation at 20,100 × g for 10 min. Gel bands were picked, proteins processed using the OMX-S protocol (Soliden, Seefeld-Hechendorf, Germany) and peptides processed and concentrated using C18 Spin Tips (PierceTM, Thermo Scientific). In brief, Spin Tips were preactivated using 20 µl 80 % (v/v) acetonitrile, and centrifuged for 2000xg, 2 min, the OMX extract was loaded and Spin Tips, centrifuged at 1200xg for 2 min. Peptides plus pigments were eluted using 100 % acetone, to yield an 80% acetone/10 mM Hepes pH 8 solution. Eluates were directly loaded and separated on C18 reversed-phase HPTLC (Merck) plates and developed in a mobile phase composed of 30 ml methanol, 20 ml acetone, 1 ml water at RT in darkness. Plates and gels were analyzed for fluorescence of Pchl(ide) and Chl(ide) at Ex 633 nm/Em >670 nm (633/680, BP30) using a Typhoon Trio (GE Healthcare, Buckingham, GB) and for fluorescence of Chl(ide) at Ex 680 nm/Em >700 nm (680/700, BP30) using an Odyssey CLx (Biotechnology – GmbH, Bad Homburg) scanner. Emission spectra were recorded at 293 K in a Fluorolog-3 spectrofluorometer (HORIBA Europe GmbH, Gotenburg, SE) with Ex 440 nm (450-W Xenon lamp). Gel bands (protein extracts corresponding to 2.5 × 10⁷ plastids) were placed between quartz plates, and spectra (550–750 nm) were baseline-corrected and normalized at 710 nm (Gilmore and Ball 2000) or ∼630 nm.

### LC-MS/MS

LC-MS/MS was performed using nanoAcquity UPLC (Waters, Wilmslow, UK) and LTQ Orbitrap Velos (Thermo Scientific, Sunnyvale, CA, U.S.A.). Eluants were nanosprayed; *m/z* values measured at 30,000 resolution. Ions (charge ≥ 2+) were fragmented. Data were processed in Protein Discoverer v2.1 (Thermo), converted to mgf, and searched via Mascot (Matrix Science; *p* < 0.05, peptide score >20). Scaffold 4.0 thresholds: 99.9% protein, 95% peptide, ≥5 unique peptides.

### Microscale thermophoresis (MST)

LIL3.2 was expressed and purified (Mork-Jansson et al. 2015a). Chlide intrinsic fluorescence (100 nM final) was monitored in reconstitution buffer (100 mM Tris pH 11, 5 mM 6-aminocaproic acid, 1 mM benzamidine, 12.5% sucrose, 2.5 mM DDM). Unlabeled LIL3.2 was titrated (1:1 dilution, 2.5 μM to 0.61 nM). After 2 h at RT, samples were loaded into Monolith NT.115 Premium Capillaries and measured (Monolith NT.115; LED 10%, MST 40%). Data were analyzed using MST software, MO.Control, and ΔF_norm_ = baseline-corrected F_norm_ [‰] (Jerabek-Willemsen et al. 2011).

## Results

### Light-Induced Fluorescence Changes in Etioplast Membrane Proteins

The binding of chlorins, Pchl and Chl, to proteins of the etioplast membrane was investigated by fluorescence analysis of protein complexes separated by electrophoresis. Etioplasts were isolated from dark-grown or from 10-s illuminated 4.5-day-old barley seedlings. Membranes were solubilized and protein complexes separated using native LDS-PAGE (1D LN-PAGE). The protein subunits were released from the native protein gel bands and separated by a low denaturing second dimension gel (2D SDS-PAGE) (Fig.1). Changes in the chlorin binding to protein were determined by fluorescence emission analysis at >670 nm upon in-gel excitation scanning of the 2D gels at 633 nm (Fig. 1).

**Figure 1:**
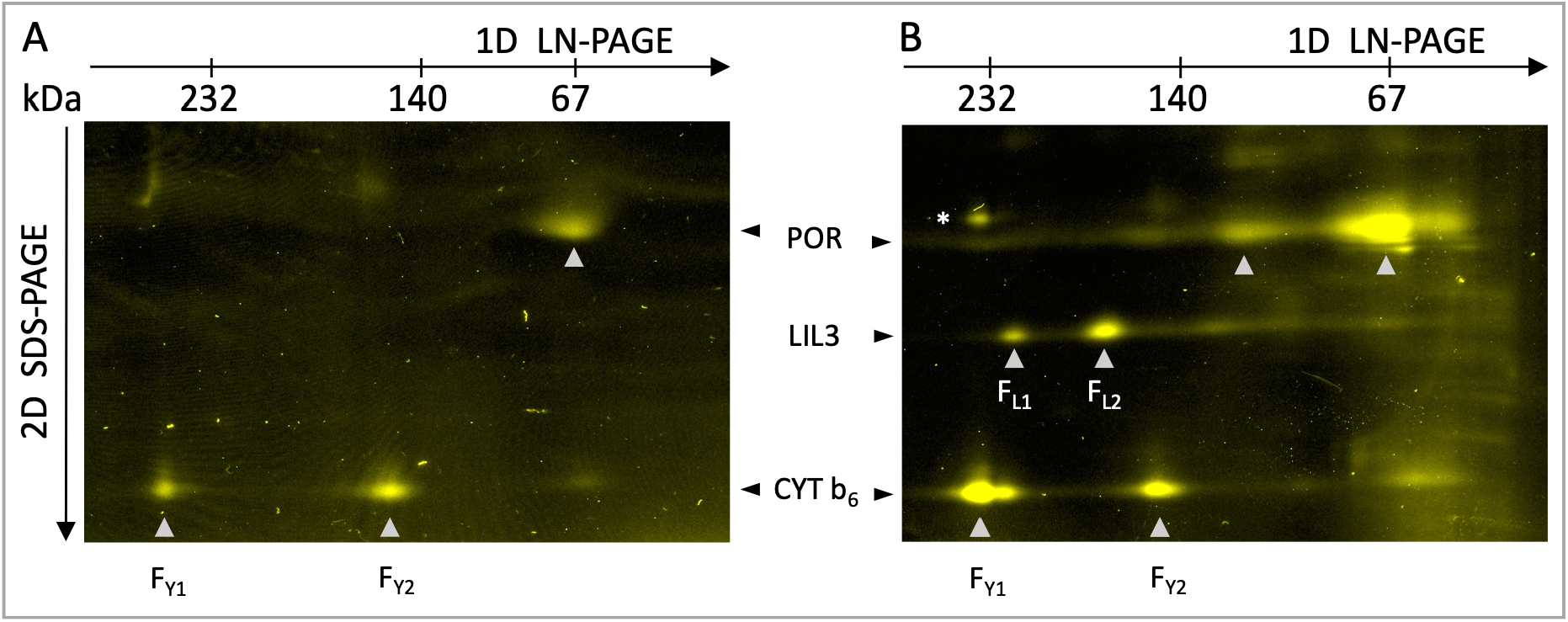
2D-PAGE analysis of light-induced fluorescence changes in etioplast membrane proteins. Etioplast membrane protein complexes were isolated from etiolated barley seedlings (A) or from etiolated plants illuminated for 10 seconds (B). Membranes from 2 × 10^8^ etioplasts were solubilized, and protein complexes were separated according to molecular weight by LDS-Native PAGE (LN-PAGE) in a first dimension (1D, horizontal arrow) and by denaturing SDS-PAGE in a second dimension (2D, vertical arrow). Protein subunits were detected in the 2D gels by in-gel excitation/emission scanning at wavelength settings of 633/670 nm, and proteins were identified as LIL3, Cyt b6, PORA, and PORB (POR) by gel blot and MS analysis. Mobility of the corresponding protein complexes Cyt b6f dimer (F_Y1_) and monomer (F_Y2_), are indicated below the 2D gel. The relative mobility of the HMW native molecular weight standards (kDa) are shown on top of the 2D gel. The mobility of the protein subunits in the 2D gel is labelled between gels A, and B. Vertical arrowheads in the gel mark the mobility of assembly states with increasing MW of POR, LIL3, and Cyt b6f complexes (A and B).

Fluorescent protein complexes in the native gels were localized at a molecular weight of about 67 kDa, and at 150 and 300 kDa. Proteins were identified as POR, and Cyt b6f in its dimeric (F_Y1_) and monomeric (F_Y2_) assembly state in the 1D gel and via the corresponding protein subunits in the 2D gel as PORA and PORB at a molecular weight of about 35 kDa (POR), and as Cyt b6 (F_Y1_, and F_Y2_) at about 19 kDa (Fig. 1A, arrows, Fig. 2A, Fig. 3A). When plants were illuminated for 10 s before isolation of the etioplast protein complexes, the 2D analysis revealed increased fluorescence intensity in POR and in the Cyt b6subunit corresponding to the dimeric Cyt b6f complex (F_Y1_) (Fig. 1). In addition, two fluorescent proteins of about 30 kDa were newly formed, which were identified as LIL3 by MS and gel-blot analysis (Fig. 1B, LIL3). The protein specific fluorescence increase indicated the light-dependent synthesis and accumulation of Chlide and/or Chl in POR, Cyt b₆, and LIL3. Also a fluorescent labeling of Cyt f was noted (Fig. 1, *). To our surprise, no Chl-binding proteins of the two photosystem complexes of PSI and PSII were identified (Fig. 1). This indicated that, upon induction of Chl synthesis in-vivo, binding of Chl to Cyt b6f during de-etiolation precedes accumulation of Chl in the reaction center photosystem complexes. We therefore investigated the Chl dependent assembly of the photosystem complexes during polysome run-off conditions in etioplasts.

**Figure 2:**
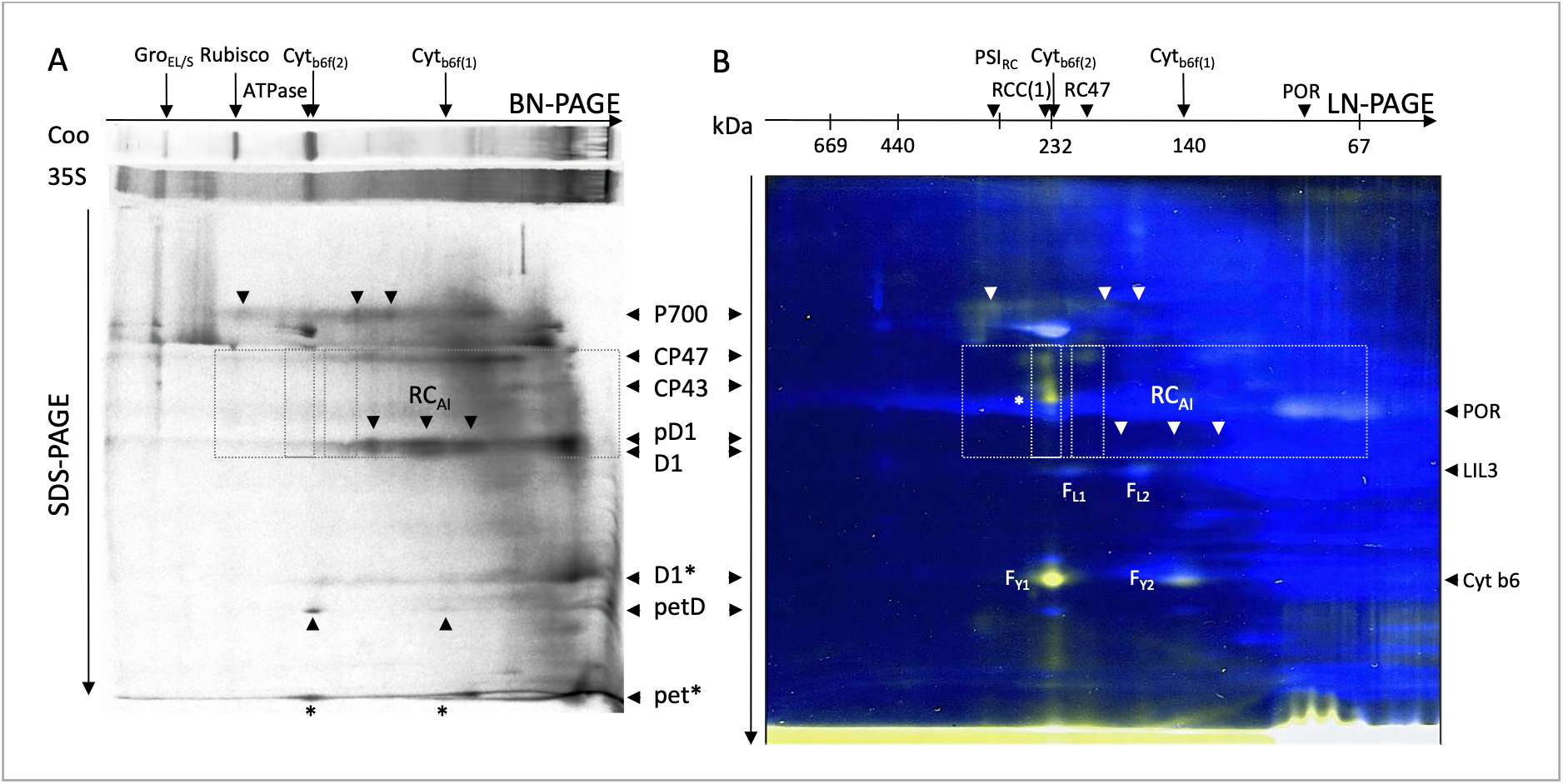
Visualization of etioplast membrane proteins by radiolabeling, autofluorescence and Cy2 labeling. Etioplast (1 x 10^8^) were either isolated from 4.5-day-old dark-grown barley seedlings (A) or after illumination of etiolated seedling for 1h (B). Etioplasts, were pulse-radiolabelled with 35S-Methionine in the dark, and radiolabel chased for 80 min in the presence of Chl synthesis (A) or etioplast membrane proteins were labelled with CyDye (Cy2) (B). Membranes were solubilized, and proteins separated by 2D Native-PAGE (BN-PAGE, A or LN-PAGE, B). Radiolabel in proteins was detected via Phosphoimaging (A), or protein specific chlorin fluorescence was recorded by scanning the gel at 670 nm upon excitation at 633 (yellow spots) plus Cy2 fluorescence at 520 nm upon excitation at 488 nm (Cy2, blue) (B). The two fluorescent protein images (B) were merged including color labeling using ImageQuant software (Methods). The mobility of radiolabelled protein subunits of PSI (P700), of PSII (CP47, CP43, pD1, D1, D1*), and of the reaction center assembly intermediate complexes (RC_AI_), and of Cyt b6f complexes (petD, pet*) are marked using in-gel vertical arrow heads. The dashed box shows the relative mobility of the radiolabeled (A) and of the Chl binding (B) protein subunits in the 2D gel which point to the location of distinct protein complexes in the 1D gel. Protein subunits, and protein complexes were identified by gel blot and/or by MS analysis. Relative mobility of the protein bands in the 1D Blue native gel (BN-PAGE) is shown as horizontal gel slabs upon staining with Coomassie G250 (Coo) or radiolabel detection (35S) (A). HMW native molecular weight standard protein kit (kDa) (GE Healthcare) is provided on top of the 1D LN-PAGE. The location of radiolabel and of Chl fluorescence associated with protein subunits of PSI (P700) and PSII (CP47, CP43), and of the Cyt b6f complexes F_Y1_, and F_Y2_ (Cyt b6, petD, pet*), and of LIL3 complexes, F_L1_, F_L2_ (LIL3) is indicated in the 2D gels. Radiolabel in pet* in Cyt b6f refers to the three plastid encoded subunits L, N, and G identified by MS. The location of protein complexes GroEL/S, Rubisco, ATPase, PSII complexes RCC(1), RC47, Cyt b6f monomer (1), and dimer (2) and of POR in the 1D gels labelled (top, horizontal arrow).

**Figure 3:**
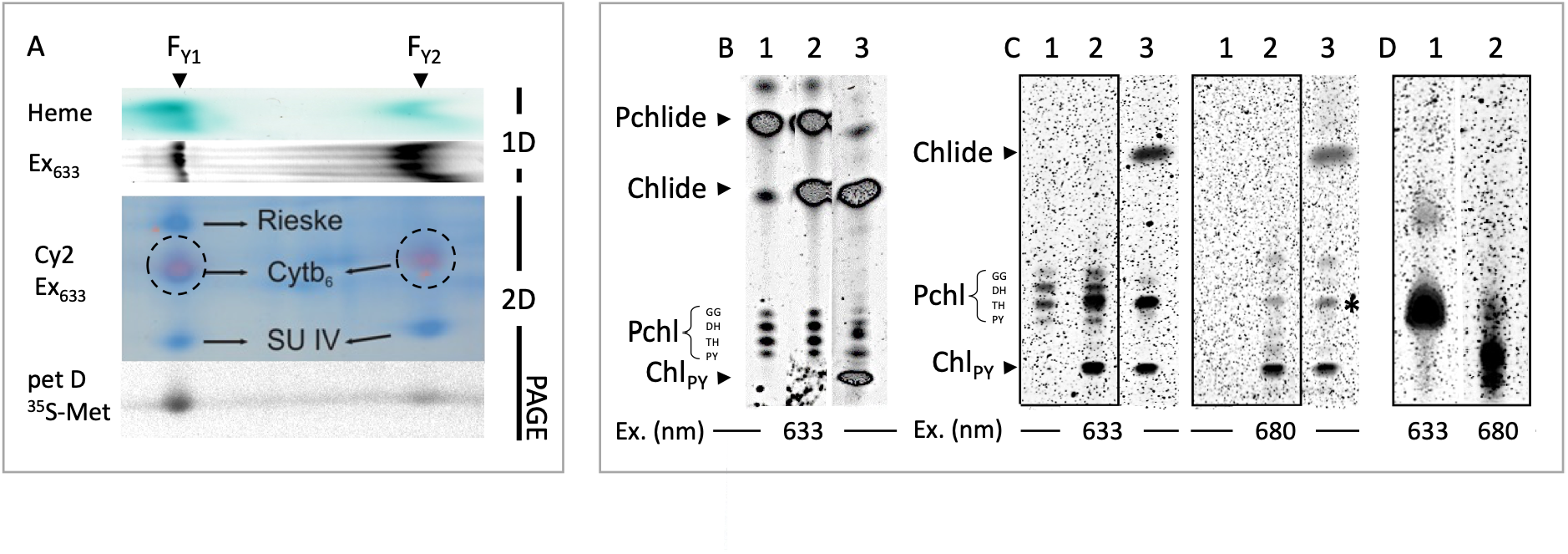
Characterization of the Cyt b6f dimer. The Cyt b6f complex was investigated in etiolated plants illuminated for 10 seconds after 2D LN-PAGE separation of etioplast membrane proteins (A, 1D, 2D PAGE). The Cyt b6f complex was identified in the 1D gel via the Cyt specific Heme stain and the Pchl/Chl fluorescence (A, Heme, Ex633). Protein subunit characterization in the 2D gel identified the Rieske-Iron sulfur (Rieske), Cytochrome b6 (Cytb6), and Subunit IV (SU IV) by MS analysis of the Cy2 stained protein spots (A). The Cyt b6 was further identified by Pchl/Chl binding (1D and 2D, Ex633), and SU IV by 35S-Methionine radiolabeling of petD gene expression (2D, petD 35S-Met). Labeling was recorded by scanning the gels by excitation/emission scanning at 633/670 nm (BP30) (Chl, violet color overlay) and at 488/520 nm (Cy2, BP40) and photostimulated luminescence (35S-Met). Chlorins were separated by HPTLC chromatography and the fluorescence of plates scanned for Pchl(ide) and Chl(ide) at 633/670 nm or Chl(ide) at 680/700 nm (B to D). Chlorins were extracted from the etioplast membranes of plants kept in darkness (B, lane 1), or illuminated for 10s (B, lane 3) or from plastids illuminated in the absence of GGPP (B, lane 2). After separation by 1D PAGE, Chlorins were extracted from the Cyt b6f complex dimer gel-band (F_Y1_) (C, lanes 1, and 2, boxed). The Cyt b6f complex was isolated from plastid membranes of plants kept in darkness (C, lanes 1) or from plants illuminated for 10s (C, lane 2), or chlorins were extracted from the vesicle front of the 1D gel (C, lanes 3). Chlorins from the Cyt b6 subunit were extracted from the fluorescent gel-spots after 2D-PAGE and visualized at 633/670 nm (D, lane 1), and 680/700 nm (D, lane 2).

### Chl binding to Cyt b6f precedes PSI and PSII assembly

Etioplasts were pulse-radiolabeled in the dark for 15 min using 35S-methionine (Suppl. Fig. 1). In the subsequent 80 min radiolabel chase, etioplasts were supplemented with L-methionine and Lingomycin and GGPP and Chl synthesis induced by illumination for 10 s. The assembly of the photosystem complexes was analysed upon separation of the solubilized protein complexes by 2D BN-PAGE (Fig. 2A).

Translation of the plastid-encoded proteins resulted in radiolabeling of protein subunits of PSI (P700), and PSII (CP47, CP43, D1), Cyt b6f (petD, pet G, L/N), Rubisco (LSU), ATPase (α-, and β-SU) (Fig. 2A). The 2D protein subunit analysis showed that holocomplexes Rubisco and the CF1 ATPase complex were radiolabeled and also, a higher-MW band accumulated both the ATPase α/β subunits and the LSU subunit in the distinct molecular weight complexes. The higher-molecular weight complex radiolabel by LSU and ATPase α/β subunits was stained with Coomassie in the 1D native gel and identified as the GroEL/S chaperon (Fig. 2A, GroEL/S). Interestingly, also radiolabel in PSII subunits CP43 and CP47 were colocalized with the GroEL/S protein band (Fig. 2A). This finding showed the principle for reading the 2D gels. The differential accumulation of radiolabeled protein subunits in the higher molecular weight regions of the 1D gel resulted in accumulation of a protein subunits in the 2D gel in a vertical line defined by the electrophoretic field of the second dimension separation. The field specific arrangement of these proteins indicated their stable interactions ie. during assembly of the subunits into the holocomplexes. In the lower molecular weight region of the 1D gel, radiolabeling of the native protein complexes was evident within the 2D gel as horizontal radiolabel extensions along the electrophoretic field of the native 1D gel. With respect to the Chl binding photosystem proteins, the PSI and PSII core proteins (P700, CP47, CP43, D1, D1*) showed accumulation in distinct protein assembly states with increasing molecular weight (Fig. 2A, downward arrow heads). The subunits of P700 showed a low quantity of radiolabel accumulation at the molecular weight of the reaction center complex and in two lower molecular weight assembly intermediates (Fig. 2, P700, arrow heads). For PSII, pD1 showed the highest radiolabel accumulation in non-assembled protein, and the pD1/D1-protein assembled in three reaction center assembly intermediate complexes (RC-AI). In parallel, radiolabel was found in non-assembled CP47 and CP43 protein and CP47 showed a molecular weight extension of its radiolabeling but without a colocalization of radiolabel in the distinct reaction center assembly intermediates (Fig. 2A, pD1, CP47, dashed squares). A similar extension relative to the CP47 protein was found for the 23 kDa breakdown product of the pD1 protein (pD1*). Results indicated for PSII that reaction center assembly was regulated independent from CP47. For the Cyt b6f complex, radiolabeled petD was very selectively accumulated in the dimeric Cyt b6f complex. Also, the small plastid encoded subunits of the Cyt b6f complex were expressed and assembled distinctly with the dimeric complex; however, resolution of the gel was too low for assigning the radiolabel specifically to subunits petG or petL/N (Fig. 2A, pet*). Results showed that with the onset of Chl synthesis, newly synthesized protein subunits of PSI and PSII assembled directly in photosystem reaction center protein complexes of PSI and PSII; however, quantities of subunits in the assembly states were low, and for PSII no assembly of CP47 or CP43 into RC47 or the mono- and di-meric reaction center core complexes could be identified (Fig. 2A, dashed squares). In contrast, Cyt b6f dimers showed no assembly intermediates besides the Cyt b6f monomer, and distinct radiolabeling allocation of petD and of petG/L/N to the dimeric Cyt b6f complex indicated that the assembly process was efficient. Results documented the progression of photosystem biogenesis in etioplasts directly upon induction of Chl synthesis, which raised the question why Chl binding was only identified for the Cyt b6f dimer.

The 2D gel of 10s illuminated plants had shown strong fluorescence increase only for the POR, LIL3, and the Cyt b6 protein subunits while fluorescence from PSI or PSII proteins was absent (Fig. 1B). We therefore extended illumination of the etiolated plants for 1 h and labeled membrane proteins using Cy2 for a dual in-gel location of Chl and protein. The in-gel detection of both Chl fluorescence and protein allowed to match the 2D protein subunit location from radiolabeling, Cy2 labeling and Chl fluorescence (Fig. 2B). In the 1h illuminated in-vivo state, the highest molecular weight Chl fluorescence overlapped with the radiolabeled position for the PSI reaction center complex determined upon in-vitro induction of Chl synthesis. Chl fluorescence in the P700 spots was also detected in the lower molecular weight extension of the radiolabeled P700 intermediates (Fig. 2A and B, black and white downward arrow head, PSI_RC_). After 1h illumination, also PSII proteins CP47, and CP43 showed accumulation of Chl in two complexes, the monomeric reaction center core (RCC(1)) and the RC47 complex while no Chl accumulation in the three distinct RC assembly (RC-AI) intermediates was found (Fig. 2B, downward arrowhead; PSII_RCC(1)_; RC47). This finding indicated the primary importance of CP47 and CP43 in PSII and the corresponding C-terminal protein sequences in the psaA, and psaB proteins to stabilize Chl binding and accumulation of photosystem complexes. In contrast, the assembly of reaction center protein of PSII indicated that here protein interactions were dominating. Data indicated that Chl binding to the reaction center structure parts was coordinated after binding of Chl to the inner antenna parts. However, even after 1 h of illumination of the plants, the overall fluorescence yield from PSI and II protein complexes was maintained very low relative to accumulation of Chl in Cyt b6f dimers (Fig. 2B, F_Y1_).

Comparison of the 10s and 1h in-vivo illuminated etiolated plants showed that Chl accumulation in Cyt b6f was several-fold stronger in both conditions relative to the accumulation of Chl in the photosystem complexes after 1 h of illumination. This indicated that Chl accumulation in PSI and PSII complexes was delayed (Fig. 1B and 2B). The specific radiolabeling accumulation of newly expressed petD and petG/L/N protein subunits in the dimeric Cyt b6f underscored the productive and dominant channeling of Chl for assembly of dimeric Cyt b6f. However, while accumulation of Chl in Cyt b6f was directly triggered by the onset of Chl synthesis, petD turnover was independent of an alteration of Chl synthesis in the in-vitro and in-vivo conditions indicating that the regulation of protein turnover and Chl binding are regulated independently (Suppl. Fig. 2).

### Characterization of Pchl and Chl binding to the Cyt b6f dimer

Although it appeared clear that the origin of the fluorescence in the two assembly states of the Cyt b6f complex was correlated with Chl synthesis we isolated the pigments directly from the Cyt b6 dimer bands to identify the pigment. The location Cyt b6f in the 1D gel was verified upon 10 illumination in-vivo. The presence of Cyt b6 and Cyt f were verified via the covalently linked hemes and the presence of Pchl or Chl correlated via the fluorescence upon excitation at 633 nm (Fig. 3A, Heme, Ex633 nm). The Cyt b6f dimer showed both the Heme stain in the Cyt b6f dimer and fluorescence and the gel bands overlapped (Fig. 3A). The 2D analysis of the two 1D gel-bands identified the Cyt b6f via the Cy2 specific staining of the subunits, Rieske-Iron-Sulfur, Cyt b6 and SUIV, and the bands were specific for the petD expression and radiolabel incorporation in the SUIV protein. In addition, the 2D location of the Cy2 label of Cyt b6 specfically overlapped with the fluorescence recorded in the dimer and monomer Cyt b6f band (Fig. 3A, and Suppl. Fig. 2).

The fluorescent pigments bound to the Cyt b6 protein in the Cyt b6f dimer complex were investigated using organic phase extracts. Etioplast membrane extracts were compared with extracts from the Cyt b6f dimer and the Cyt b6 subunit gel bands (Fig. 2B to D). Extracted pigments were separated by high-performance thin-layer chromatography (HPTLC) and pigments distinguished by their chromatographic mobility and fluorescence upon excitation at 633 and 680 nm (Methods). Isolated etioplast membranes were found to contain Pchlide, a low amount of Chlide, and Pchl in the four distinct hydrogenation states (Fig. 2B, lane 1). Illumination of isolated etioplasts in the absence of supplementation of etioplasts with GGPP, triggered synthesis of Chlide, however, the content of Pchlide appeared less affected and no Chl accumulation was found (Fig. 2B, lane 2). For induction of Chl synthesis, etiolated seedlings were illuminated for 10 s, then etioplast membranes were prepared on ice and pigments were extracted. Under this condition, Pchlide was strongly diminished, synthesis of Chlide was induced, the content of all Pchl bands decreased and Chl accumulated in the phytylated state (Fig. 2B, lane 3). In addition an additional Chl band with a Pchl-like mobility was noticed (Fig. 2B, lane 3, *). When pigments were extracted from the Cyt b6f complex isolated from the etioplast membranes of plants either maintained in darkness or illuminated for 10s of the plant and fluorescence in the HPTLC plates analysed upon excitation at 633 nm, we identified Pchl in the absence (Fig. 2C, lane 1, boxed) and Pchl plus Chl in the presence of Chl synthesis (Fig. 2C, lane 2, boxed). All four hydrogenation forms of the geranylgeraniol side chain of Pchl extracted from etioplast membranes (Fig. 2B, lane 1) were also extracted from the Cyt b6f dimer (Fig. 2C, lane 1); however, the Cyt b6f complex did not bind Pchlide or Chlide. Fluorescent pigments from the 10s illuminated plants that were not bound to protein were located at the gel-front after 1D PAGE (Fig. 3C, lane 3). Here, Chlide, phytylated Chl_PY_ and the Chl derivative were present (Fig. 2C, lane 3*). The binding of Chl_PY_ and of the Chl derivative was further investigated by a Chl specific excitation of the HPTLC plates at 680 nm and recording of fluorescence larger than 700 nm (Fig. 3C, lanes 1 and 2, boxed). Pchl specific bands recorded at 633 nm were now no longer detected, and also fluorescence from the Chl derivative was markedly reduced (Fig. 3C, lanes 1 and 2, 680 nm). The selective excitation of Chlide/Chl was also confirmed in the pigments extracted from the gel-front (Fig. 3C, lane 3, 680 nm). We also extracted the chlorins bound to the Cyt b6 protein subunit after 2D-PAGE separation (Fig. 3D). The fluorescence analysis confirmed Pchl and Chl binding at 633 and 680 nm excitation (Fig. 3D). The pigment extractions from the Cyt b6f gel-bands corroborated that esterified Chl_PY_ and a Chl derivative were extracted from the etioplast membrane and the Cyt b6f dimer and that the Cyt b6f complex deselected non-esterified chlorins. The specific and rapid binding of Chl to the Cyt b6f dimer allowed us to control its Chl specific biogenesis by controlling access to Chl in-vitro.

### Temperature-dependent Chl accumulation in Cyt b6f monomers vs. dimers

The kinetics and synthesis of Chl can be controlled in isolated etioplasts by the reaction temperature via the enzyme chlorophyll synthase (EC 2.5.1.62) (CHS). In contrast, substrates Chlide and GGPP or PhPP, require supplementation either by photo-transformation of Pchlide which triggers Chlide synthesis, or Chlide supplementation, while Chlide esterification to Chl requires supplementation of GGPP or PhPP. Here, the binding of Chl to the protein complexes of the etioplast membrane was analyzed 0, 10 and 20 min after a 10-s illumination by 1D LN-PAGE (Fig. 4). In the non-illuminated etioplast control (E), both fluorescent dimeric and monomeric Cyt b6f complexes, F_Y1_ and F_Y2_ were determined, respectively (Fig. 4A, lane E, F_Y1_ and F_Y2_).

**Figure 4:**
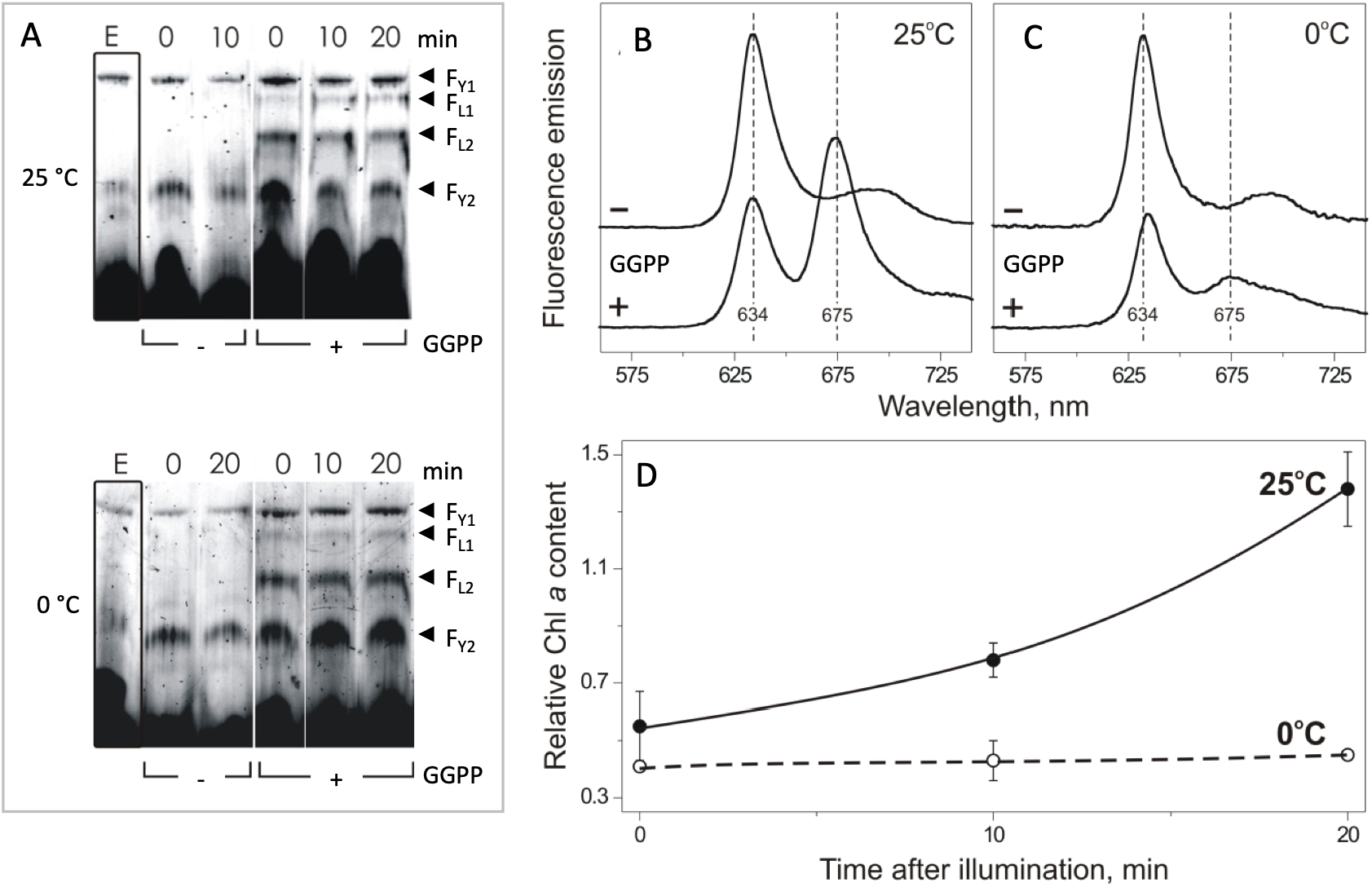
Temperature-dependent accumulation kinetic of Chl in the Cyt b6f dimer. The kinetics of Chl accumulation in dimeric (F_Y1_) and monomeric (F_Y2_) Cyt b6f gel-bands was investigated in illuminated etioplasts (5 × 10^7^ plastids). Membranes were solubilized after 0, 10, and 20 min of illumination in the absence (minus) and presence (plus) of 2.77 nmol GGPP. Protein complexes were localized by fluorescence scanning after separation by LN-PAGE (A). Fluorescent gel-bands were labelled according to gel-blot analysis using antibodies against Cyt b6 (F_Y1_, F_Y2_) and LIL3 (F_L1_, F_L2_). Kinetic changes in the fluorescence intensity at 670 nm were determined upon excitation at 633 nm after a reaction time of 0, 10, and 20 min after illumination of etioplasts in the absence (minus) and presence (plus) of GGPP (A). The fluorescence emission spectra were recorded for dimeric Cyt b6f band, F_Y1_, after a 20 min reaction time at 0 °C and 25 °C from 560 to 740 nm with excitation at 440 nm, and spectra were normalized to the mean fluorescence intensity at 710 nm (B, C). The change in the relative Chl content in the F_Y1_ gel-band was determined as the 675 to 634 nm ratio of the fluorescence emission maxima during the 20 min reaction time (D).

When complexes were directly isolated after illumination of etioplasts in the absence of GGPP (Fig. 4A, lane 0, minus GGPP), fluorescence yield >670 nm from the F_Y2_-band increased upon excitation scanning at 633 nm relative to the complexes isolated without illumination (Fig. 4A, lane E, minus GGPP). In contrast, fluorescence yield from the F_Y1_-bands remained about constant (Fig. 4A, lane 0, minus GGPP). However, when etioplasts were illuminated and thereafter were incubated in darkness for an additional 10 to 20 min, before complex isolation, the signal from both assembly states decreased (Fig. 4A, lanes 10/20, minus GGPP). This result was very similar in etioplasts maintained at 25 and 0 °C. The fluorescence increase indicated that monomeric, but not dimeric, Cyt b6f complexes were binding Chlide at 25 and 0 °C, and that the Chlide binding was not kinetically stable. In contrast, etioplasts illuminated in the presence of GGPP showed an immediate and much stronger fluorescence increase in the monomeric F_Y2_-band relative to the absence of GGPP, and the dimeric F_Y1_-band also increased in fluorescence (Fig. 4A, lane 0, plus GGPP). This finding was observed for both etioplasts illuminated at 25 and 0 °C (Fig. 4A, B). For etioplasts illuminated in the presence of GGPP, the strongest change in fluorescence was noted by varying the temperature. When the dark incubation time was extended to 10 and 20 min, fluorescence of the monomeric F_Y2_-band decreased after the 0-time point at 25 °C; while at 0 °C, fluorescence increased in the 10-min incubation time point and remained elevated at the 20-min incubation (Fig. 4A, B, 0 to 20 min, plus GGPP). In contrast, fluorescence recorded from the F_Y1_-band remained about constant throughout the incubation time in the presence of GGPP at 25 and 0 °C (Fig. 4A, B, F_Y1_). In addition to the change in the fluorescence of the Cyt b6f bands, synthesis of Chl selectively induced a change in fluorescence associated with the assembly of two LIL3 complexes (F_L1_, F_L2_) (Fig. 4A, B, plus GGPP). Especially, the kinetic changes in the fluorescence yield of the LIL3 F_L2_-band mirrored the changes of the F_Y2_-band at both 25 and 0 °C. The synthesis of Chlide in the absence of GGPP did not trigger an accumulation of a fluorescent LIL3 complex at both temperatures (Fig. 4A, B, minus GGPP). The Chl-dependent fluorescence yield in the F_L2_-bands was found increased relative to the F_L1_-band at both 25 and 0 °C. However, at 0 °C the signal yield was maintained but decreased at 25 °C indicating lower stability of Chl binding in this assembly state at 25 °C (Fig. 4A, B, F_L1_, F_L2_).

For the monomeric Cyt b6f F_Y2_ -band, the differential changes in the fluorescence yield indicated that Chlide and/or Chl synthesis and binding could have increased more rapidly at 25 than at 0 °C (Fig. 3, lane 0, plus GGPP) or that the binding was temperature-dependent with a higher stability at 0 °C (Fig. 4A, B, lanes 0 to 20 min, plus GGPP). We speculated that a selective decrease in the signal strength of the monomeric F_Y2_-band at 25 °C could indicate its dimerization, which was halted at 0 °C when the signal strength of the F_Y2_-band was maintained throughout the incubation time. However, the fluorescent gel scans did not show an increase of Chl fluorescence in the dimeric F_Y1_-band at 25 °C. Alternatively, a slower rate of Chl synthesis at 0 °C could cause a slower Chl accumulation in the monomeric Cyt b6f F_Y2_-band while the increased signal could represent the accumulation of unesterified Chlide in the band. In addition, changes of the in-gel fluorescence in the F_Y1_-band could be related to the fixed excitation wavelength at 633 nm which provided a more effective basis for fluorescence yield from the Pchl-Cyt b6f F_Y1_-band relative to the Chl-Cyt b6f complexes especially if the accumulation of Chl-Cyt b6f dimers was small within the F_Y1_-band. We investigated the accumulation of Chl in the F_Y1_-band first spectrophotometrically (Fig. 3B– D).

### Accumulation of Chlide/Chl in the F_Y2_ band precedes the F_Y1_ Cyt b6f band in-vitro

The spectral fluorescence changes of the dimeric F_Y1_-band were investigated 20 min after photo-transformation of Pchlide and induction of Chlide and Chl synthesis at 0 and 25 °C (Fig. 3B–D). The F_Y1_ native gel bands were excited at 440 nm and fluorescence emission spectra recorded from 560–740 nm (Fig. 3B, C). Fluorescence of Chl with a maximum at 675 nm was only recorded for the 25 °C incubation of etioplasts in the presence of GGPP (+) (Fig. 3B, spectrum plus). The absence of Chl accumulation (675 nm) at 0 °C in the presence of GGPP confirmed the 25 °C temperature requirement for accumulation of Chl in the F_Y1_-band. The emission maximum of 634 nm showed that at both temperatures a higher amount of Pchl was recorded in the absence of GGPP than in its presence (Fig. 3B, spectrum minus vs plus). This temperature-specific accumulation kinetics of Chl relative to Pchl was best shown by utilizing the ratio between the Chl/Pchl spectral peak areas (675/634 nm) (Fig. 3D). At 25 °C, the relative Chl to Pchl content in the dimeric F_Y1_-band increased about 2.5-fold within the 20-min incubation time, but remained unchanged at 0 °C. This substantiated our finding that phototransformation of Pchlide to Chlide progressed at 0 °C while Chl accumulation and dimerization of the monomeric Cyt b6f complexes progressed effectively only at 25 °C, leading to selective accumulation of Chl in the F_Y1_-band at 25 °C. In contrast, the Chlide-specific changes of the fluorescent yield in the F_Y2_ band were in conflict with the finding that neither Chlide nor Pchlide were found bound to the Cyt b6f dimer which indicated that a binding of Chlide to the Cyt b6f monomer was not sufficient to trigger dimerization. Alternatively, accumulation of Chlide could indicate binding to another protein in the F_Y2_ band (Fig. 4A–D).

### Chlide and Chl trigger LIL3 to comigrate with the Cyt b6f monomer and dimer

We investigated the specific effect of Chlide on Cyt b6f assembly by supplemention of etioplast membranes with external Chlide in the dark (Fig. 5A, lane 5) and by synthesis of Chlide via photo-transformation of Pchlide (Fig. 5, lane 3). Both reactions were conducted in the absence of GGPP supplementation, and Chlide-specific binding to the protein complexes was investigated by native gel separation of the complexes. To our surprise, supplementation of etioplast membranes with Chlide in the absence of light and GGPP supplementation resulted in a selective fluorescent band at the molecular weight of the monomeric Cyt b6f related F_Y2_ band, whereas no change in the fluorescence at the molecular weight of the dimeric Cyt b6f F_Y1_ band was recorded in the in-gel scans relative to the dark control (Fig. 5A, lane 1 vs. 3 and 5). A parallel gel-blot analysis of the native gel for the Cyt b6 and LIL3 protein confirmed that both Cyt b6 and LIL3 were present in the F_Y2_ band in the presence of Chlide while only Cyt b6 was also present in the F_Y2_ band in the dark control (Fig. 5A and B, F_Y2_ and C, band 2). This indicated that the Chlide specific fluorescence yield in the F_Y2_ band relative to the dark control was LIL3 specific (Fig. 5A, lanes 1 vs 3, and 5).

**Figure 5:**
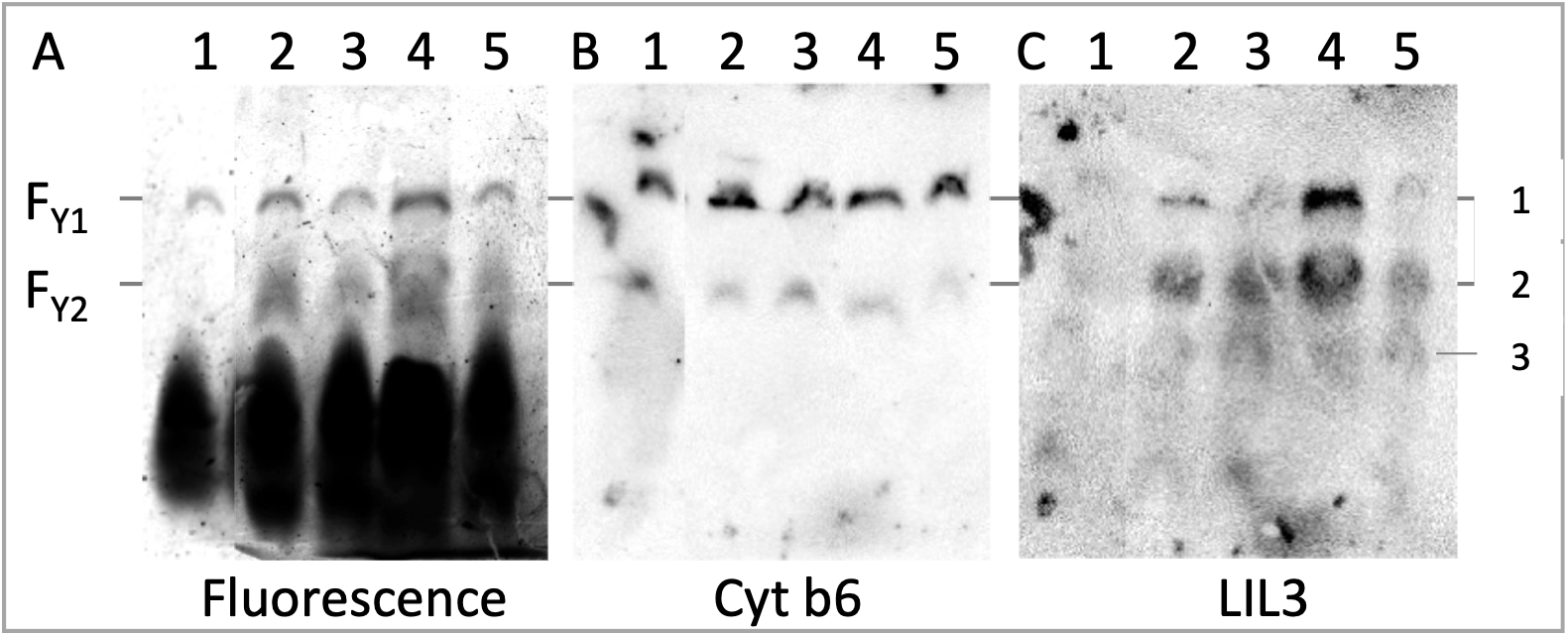
Chl synthesis triggers co-mobility of LIL3 with Cyt b6f complexes. Isolated etioplast membranes (10^7^) were incubated for 1 minute in the presence of NADPH (5 mM) in darkness (lane 1), and supplemented with GGPP (lane 2), illuminated for 10s (lane 3), supplemented with Chlide plus 2.8 nmol GGPP (lane 4), or with Chlide (lane 5). Etioplast membranes were solubilized, and protein complexes were separated by LN-PAGE. Gels were scanned for fluorescence using an excitation/emission wavelength of 633/670 nm (A, Fluorescence). Protein complexes were identified by gel-blot analysis using antibodies directed against protein subunits Cyt b6 (1:10000) (B) and LIL3 (1:7500) C) (Agrisera, Vännäs, Sweden).

Most importantly, the mobility of LIL3 markedly changed when etioplast membranes were supplemented with GGPP. When Chl synthesis was induced, LIL3 selectively accumulated in the dimeric Cyt b6f F_Y1_ band, and also fluorescence in the F_Y1_ band selectively increased while the content of the Cyt b6 in the dimer band remained constant (Fig. 5A,B,C lanes 2, 4 (Chl) vs 3, 5 (Chlide)). In addition, Chl synthesis increased the content of LIL3 in the F_Y2_ band (Fig. 5A, C lanes 2, 4). The Chlide-, F_Y2_, and Chl-specific, F_Y1_, band shifts in LIL3 mobility indicated that LIL3 shifted its associtiation from a protein complex with mobility equal to the Cyt b6f monomer in the presence of Chlide to the mobility of the Cyt b6f dimer in the presence of Chl (Fig. 5A,B, C, lane 3, 5 vs 2, 4). We therefore tested whether the amount of Cyt b6f complex would change kinetically after 1 and 10 min when Chlide/Chl was synthesized by photo-transformation of Pchlide in the absence and presence of GGPP and addressed the Chl specific change in fluorescence by isolation of the chlorins from the dimeric Cyt b6f complex band (Fig. 6).

**Figure 6:**
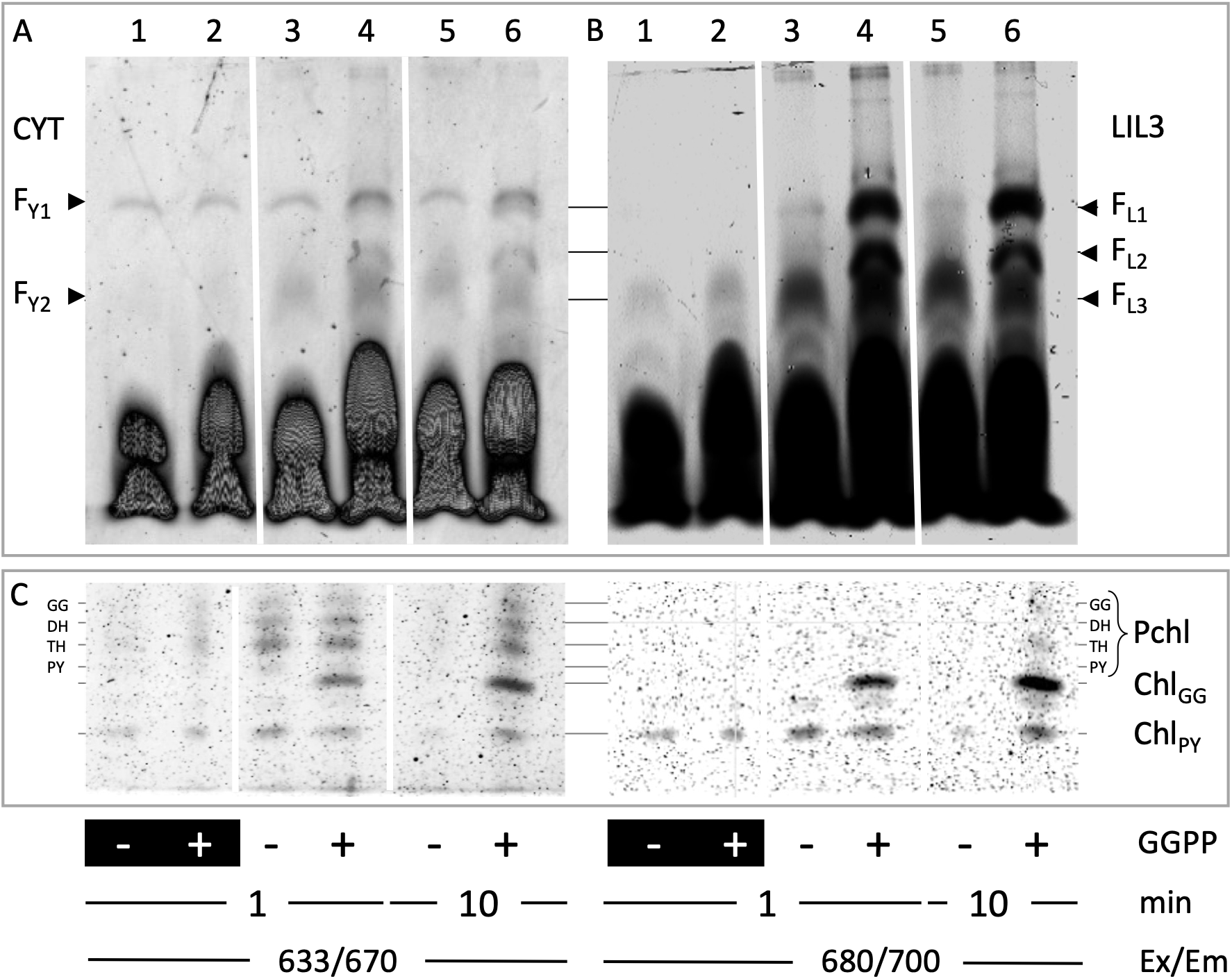
Accumulation of Chlide and Chl in LIL3 and Cyt b6f complexes. Etioplast membranes were incubated in the absence (minus, lane 1, 3, 5) and presence of GGPP (plus, lane 2, 4, 6) in the dark (lanes 1, 2) and light (lanes, 3-6). Membranes were solubilized from the membrane after a reaction time of 1 (lanes 1 – 4) and 10 minutes (lanes 5, 6) in darkness and protein complexes were separated by LN-PAGE (A, B). Gels were scanned for fluorescence with the excitation/emission (Ex/Em) wavelength set at 633/670 nm for detection of Pchl, and Chl (A) and at 680/700 nm for Chl detection (B). The mobility of fluorescent Cyt b6f protein complex bands F_Y1_, and F_Y2_ are labeled on the left and of fluorescent LIL3 protein complexes F_L1_, F_L2_, F_L3_ on the right side of the gel scans (A, 633/670, left; 680/700, right) was labelled according to gel-blot analysis using Cyt b6, and LIL3 antibodies (Fig. 4). The chlorins bound to the protein in gel-bands F_Y1_/F_L1_ were extracted from the 1D LN-gel, separated by HPTLC, and plates scanned for fluorescence emission with the same settings used for the native gels (methods) (C).

We incubated etioplast membranes in the absence and presence of GGPP (Fig. 6, -, + GGPP) for direct comparison of the effect of Chl synthesis induction. We investigated the presence of Pchl and Chl in the dimeric Cyt b6f via the 633-nm excitation and verified the presence of Chl in the Cyt b6f via the 680-nm excitation in the native gel, and upon extraction of the chlorins from the dimeric Cyt b6f F_Y1_ band (Fig. 6A, B, and C at 633/670 and 680/700 nm Ex/Em). Direct comparison between etioplasts incubated for 1 min in the absence and presence of GGPP, and in the absence and presence of light, corrobrated the Chlide specific fluorescence increase in the F_L3/Y1_ band (Fig. 6, lanes 1 vs. 3). In the presence of light plus GGPP supplementation, the fluorescence in F_L3/Y1_ shifted to the F_L2_ band, plus the highest fluorescence was now accumulating in the F_L1/Y1_ band (Fig. 6A/B, lanes 3 vs. 4). Chlorin extractions from the F_Y1_ band showed esterified Pchl in the four hydrogenation states (Pchl) and high Chl_GG_ accumulation via the 680 nm excitation scanning to be specific in the presence of GGPP supplementation. Also a low Chl_PY_ background was detected in all lanes (Fig. 6C, lanes 3 vs. 4). The analysis allowed to differentiate between the Chlide specific increase of fluorescence in the LIL3 band F_L3_ and demonstrated the Chl_GG_ specific increase in fluorescence in the LIL3 bands F_L2_ and the F_L1_ and the F_Y1_ band and results were verified in the 10 min exposures to GGPP supplementation (Fig. 6A,B,C, lanes 3/4 vs 5/6). Results demonstrated the Chlide and Chl specific accumulation and mobility shift of LIL3 during accumulation of Chl in the Cyt b6f dimer band.

Results confirmed that the induction of Chl synthesis triggered accumulation of Chl and fluorescence in LIL3 bands F_L1_ and F_L2_, and in the dimeric Cyt b6f complex band F_Y1_, and the Chlide specific fluorescence accumulation in the F_L3_ band. Based on the sequential of enzymatic steps in Chl synthesis, the shift in protein band mobility indicated that Chlide binding to LIL3 in the band F_L3_ is the origin for Chl binding to LIL3 bands F_L2_, and F_L1_, and its accumulation in the dimeric Cyt b6f band F_Y1_. The selective binding of Chlide to LIL3 could be the initial step for its chaperone activity which makes Chl synthesis and transfer to the monomeric Cyt b6f complex and its dimerization effective. We therefore investigated the direct interaction between Chlide and LIL3 in-vitro (Fig. 7).

**Figure 7:**
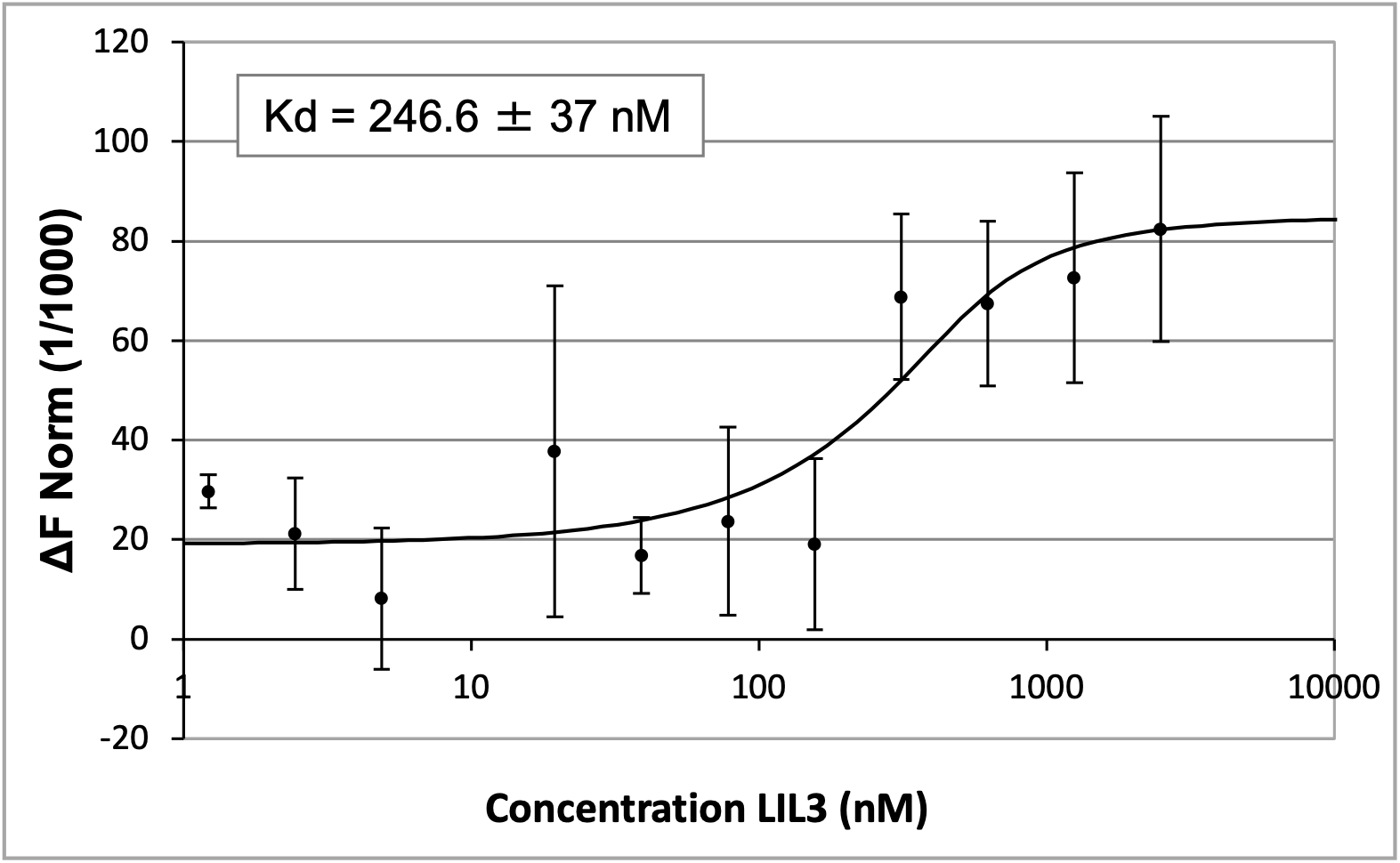
Determination of Chlide a binding to the LIL3 protein. The interaction of LIL3 with Chlide was investigated using microscale thermophoresis (Nanotemper, Munich). A constant concentration of fluorescent Chlide (0.1 μM) was incubated with increasing concentrations of unlabeled LIL3 in the presence of 2.5 mM DDM. The delta normalized fluorescence change (ΔF Norm (1/1000)) was plotted against the LIL3 concentration (B).

### LIL3 directly binds Chlide in-vitro

For investigation of Chlide binding to LIL3, Chlide was isolated from etiolated plants and the *Lil3.2* gene was isolated and cloned from *Arabidopsis thaliana*. LIL3 protein (LIL3.2) was over-expressed in *E. coli* and purified from inclusion bodies. Chlide was isolated by chromatography from etiolated barley leaves after illumination. The interaction was investigated by dissolution of an increasing amount of unlabeled LIL3.2 protein in 2.5 mM DDM in a constant concentration of fluorescent Chlide (0.1 μM) (Fig. 7). The differential exposure of the non-fluorescent LIL3 protein showed a specific change in normalized Chlide fluorescence indicating a dissociation constant for interaction with LIL3.2 of 246.6 ± 37 nM. The delta normalized fluorescence change provided evidence for a strong affinity between LIL3 and Chlide in-vitro and supported a Chlide-specific binding to LIL3 in etioplast membranes. The binding constant aligned with recent studies on pigment-protein interactions, putting weight on our finding of a specific role of LIL3 in Chlide binding and delivery during the early phase of de-etiolation studied here. This finding prompted us to differentiate between the binding of Chlide and Chl during assembly of the Cyt b6f complexes by circumventing chlorin synthesis and directly supplementing etioplast membranes with exogenous Chl *a* (Chl).

### Direct binding of exogenous Chl_PY_ to Cyt b6f

A direct binding of Chl to Cyt b6f was investigated by comparison of etioplast membranes incubated in darkness to membranes directly supplemented with 0.8 nmol phytylated Chl dissolved in 80% aceton (Fig. 8A, lane1 and 2). The fluorescence of membrane protein complexes was investigated upon separation of the solubilized complexes by LN-PAGE and native gels were scanned for Pchl/Chl excitation at 633 nm and at the Chl specific excitation at 680 nm. The binding of phytylated Chl to the complex was tested via extraction of pigments from the dimeric Cyt b6f band F_Y1_. Results showed that the fluorescence recorded at 633 nm originated mostly from the Pchl binding to the Cyt b6f dimer. The specific contribution from a de-novo binding of phytylated Chl relative to the complexes binding Pchl was visible in the increased fluorescence in both excitation scans of the 1D-gel (Fig. 8A and C, lane 1 and 2). Especially the Chl specific excitation analysis at 680 nm confirmed the specific binding of the phytylated Chl to the dimeric Cyt b6f complex in the 1D-gel scan and in the HPTLC Chl extraction analysis (Fig. 8A and C, lane 3 vs 4). The specificity of the fluorescence changes directed to the binding of Chl to the Cyt b6f complex was further corroborated by parallel gel-blot analysis showing the specific accumulation of Chl in the dimeric Cyt b6f complex. The Cyt b6 antibody showed that an about equal content of Cyt b6f complex could be recovered from the membranes in the absence and presence of exogenous Chl exposure (Fig. 8B). The in-vitro incubations demonstrated the binding of the externally supplemented phytylated Chl by the dimeric Cyt b6f complex. The Chl specific fluorescence yield was low relative to the GGPP dependent in-vitro synthesis of Chl. The lower uptake of Chl was supporting our conclusion that LIL3 might operate as a Chlid/Chl chaperone for efficient synthesis of phytylated Chl and its delivery to the Cyt b6f complex.

**Figure 8:**
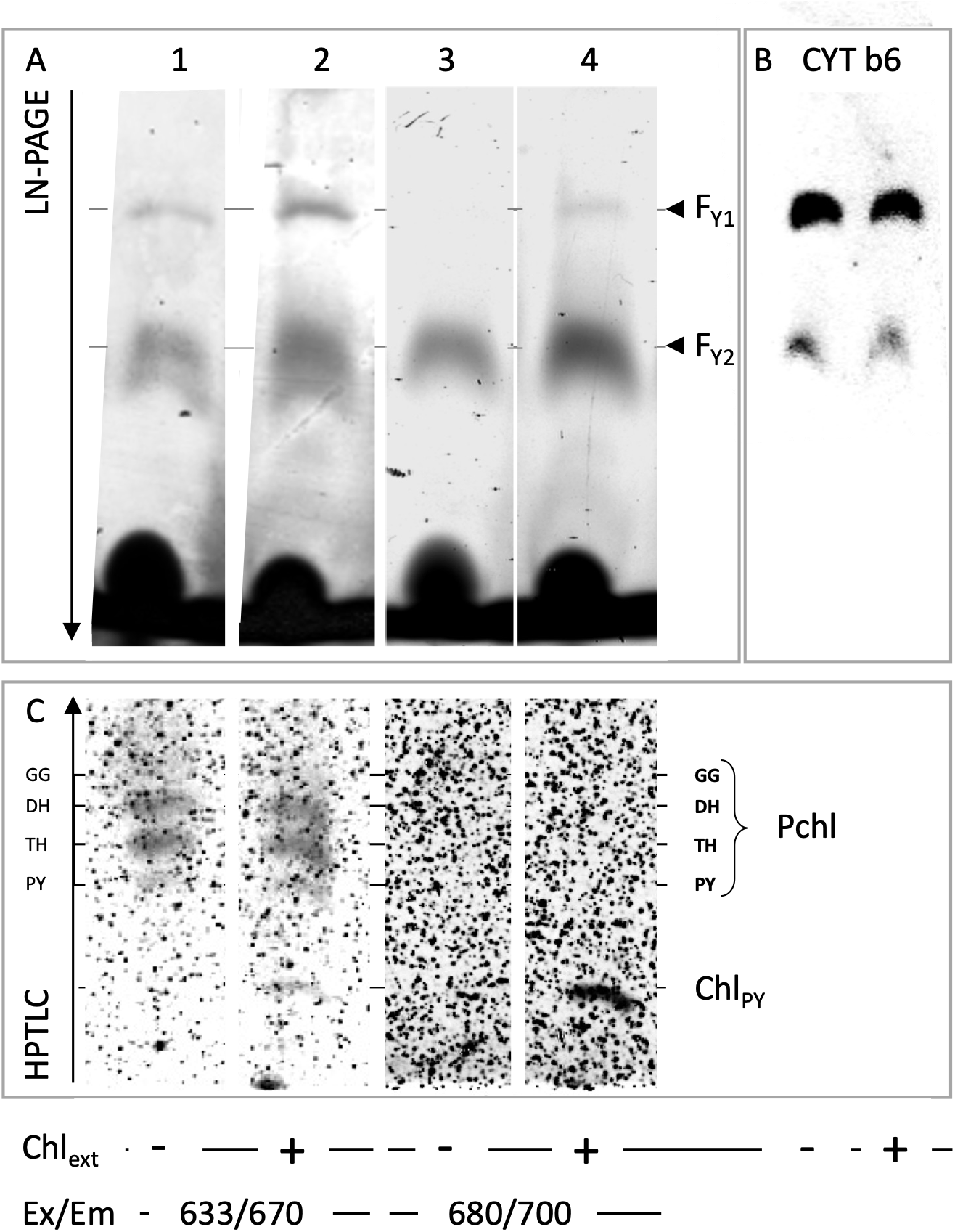
Accumulation of Chl a in the dimeric Cyt b6f complex. Etioplasts (10^8^) were resuspended in 50 mM Hepes pH 8 and 5 mM NADPH and supplemented with 0.7 nmol Chl for 1 min (Chl_ext_, lanes 2, 4), membranes were recovered and solubilized (Methods). Etioplast membranes corresponding to 2.85 *10^7^ plastids were separated by LN-PAGE. The gel was scanned for fluorescence with the excitation/emission wavelength set at 633/670 nm for Pchl plus Chl (A, lanes 1, 2), and at 680/700 nm for Chl detection (A, lanes 3, 4). The position of fluorescent dimeric (F_Y1_) and monomeric (F_Y2_) Cyt b6f complexes in the native gel are marked (A) according to gel-blot analysis using a Cyt b6 antibody (B). Pigments bound in the band F_Y1_ were extracted from the gel, separated by HPTLC, and plates scanned for fluorescence emission with the same settings used for the native gels (methods) C).

### Chl supplied via membrane lipids accumulates in Cyt b6f

We finally tested the kinetics of the Chl binding to the Cyt b6f dimer by exposure of etioplast membranes to 0.8 nmol phytylated Chl dissolved in etioplast membrane lipids. Proteins were removed from 80% acetone extracts of etioplast membranes by centrifugation and the lipid/pigment extract was supplemented with Chl_PY_ and the mixture dried by removal of the organic solvent. The extraction of Chl by the protein complexes of the etioplast membrane was then investigated by dissolution of the dried lipid/Chl mixture by native etioplast membranes (Fig. 9). The uptake kinetics confirmed the binding of Chl to the monomeric and the dimeric Cyt b6f complex and Chl fluorescence yield increased in both Cyt b6f gel bands. Spectral analysis of the accumulation kinetic was conducted upon normalization of the Chl absorbance maximum to the Pchl specific absorbance. A rapid accumulation of Chl in both the F_Y1_ and F_Y2_ bands was found at 674 nm and 676 nm relative to the Pchl signal at 632 nm and 636 nm, respectively (Fig. 9A, and C). Chl accumulation in the F_Y1_ and the F_Y2_-band increased about 5-fold relative to the background spectral levels; however, while Chl accumulated in the F_Y1_-band throughout the experimental kinetic, fluorescence in F_Y2_ showed a rapid increase followed by a decrease of the relative Chl content. Native gel analysis showed no indication for a stimulation of Chl binding to the LIL3 protein complexes throughout the kinetic supporting our conclusion that Chlide binding of LIL3 was paramount for the LIL3 specific assembly changes found in our native PAGE analysis (Fig. 9A).

**Figure 9:**
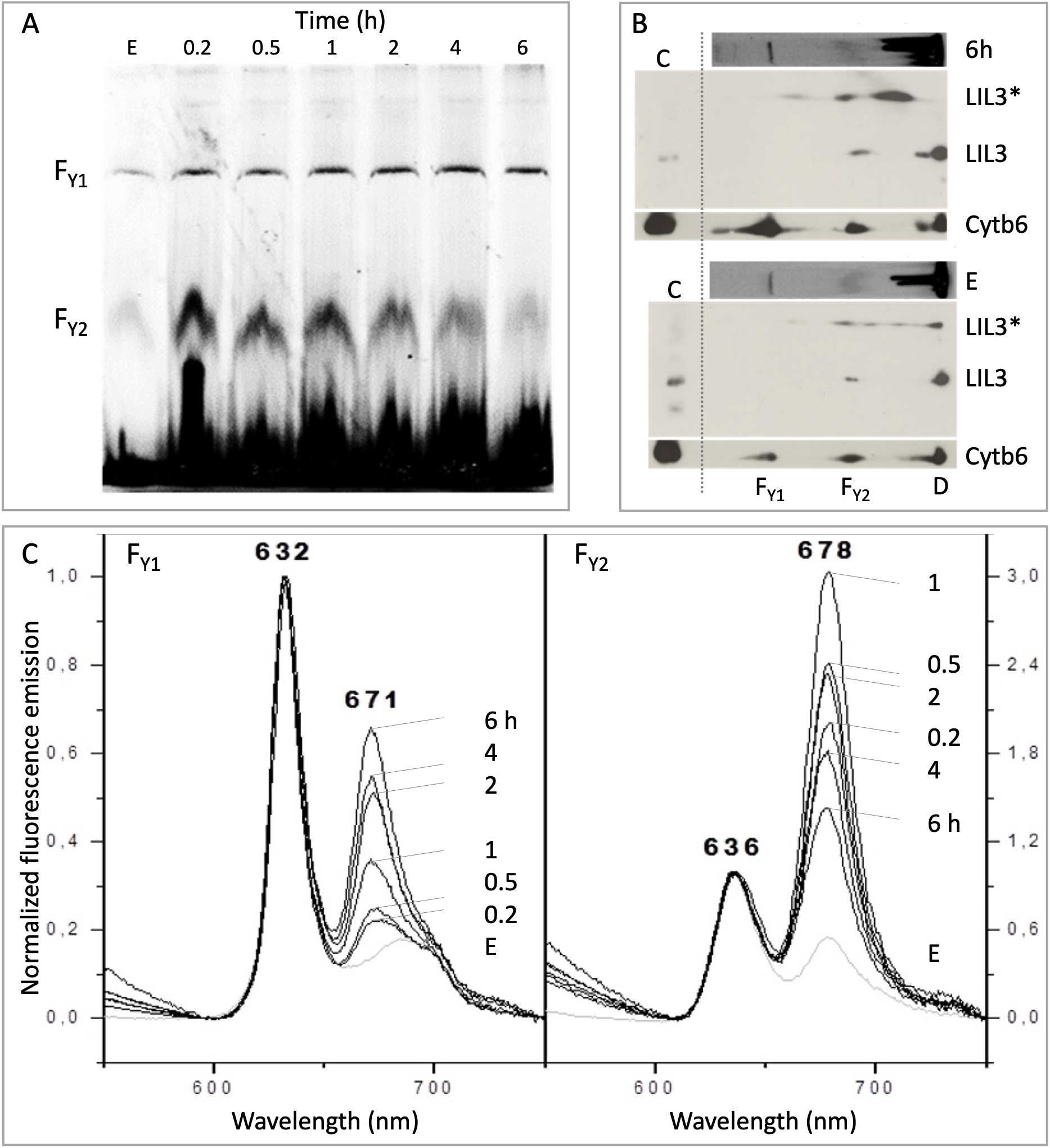
Accumulation of externally supplied Chl in the Cyt b6f dimer. Etioplasts were isolated from dark-grown barley seedlings in green safe light. Intact plastids (10^8^) were incubated with 0.8 nmol Chl for 10, 30, 60, 120, 240, and 360 min (Methods). Protein complexes of 2.6 · 10^7^ plastids were separated by 1D LN-PAGE, and the in-gel fluorescence was read out in a Thyphoon (Ex/Em, 633/670 nm) (A). The 1D lanes E, and 6 (A), were separated by a second dimension, 2D PAGE and the localization of LIL3 and Cyt b6 protein determined by gel-blot analysis (B). A denatured etioplast membrane extract from 10 s-in-vivo illuminated barley seedlings was used as a positive control for the western blot localization of Cyt b6 and LIL3 in the 2D PAGE (B, lane C). The mobility of an assembly complex of LIL3* with double the molecular weight of LIL3 was determined in the 2D PAGE analysis of Chl (B, LIL3*). For spectroscopic analysis, the gel bands at positions F_Y1_ and F_Y2_ were cut from the gel at the different time points (0 - 6 h), and the fluorescence emission spectra were recorded from 590-750 nm using 440 nm for excitation (Methods). Spectra were normalized at 632 nm for gel bands F_Y1_, and at 635 nm for gel bands F_Y2_ (C).

Nevertheless, gel-blot analysis of LIL3 and Cyt b6 protein distribution upon 2D-PAGE revealed the presence and overlap of LIL3 and Cyt b6 with the monomeric Cyt b6f F_Y2_ band (Fig. 9B). Interestingly, LIL3 was found in two different molecular weight forms after 2D-PAGE indicating its presence in a monomer and dimer state. However, in the absence of a GGPP dependent in-vitro synthesis of Chl from Chlide, no shift in the LIL3 mobility was observed upon external supplementation of phytylated Chl (Fig. 9B, LIL3*). This further supported our finding that the LIL3, L_F3_-band chaperoned Chlide binding while higher molecular weigh LIL3 L_F2_, and L_F1_ complexes chaperoned Chl esterification by CHS, and reduction of the geranyl-geraniol side chain by GGR (Fig. 10). The yield of Cyt b6 in the F_Y1_ band was increased in the gel-blot analysis indicating that the long-term Chl binding exposure supported an increase in the accumulation of dimeric Cyt b6f complex. Results corroborated our results from in-vitro synthesis of Chl and supported our conclusion that LIL3 chaperones the synthesis and effective distribution of Chl for binding to the Cyt b6f complex (Fig. 10).

**Figure 10:**
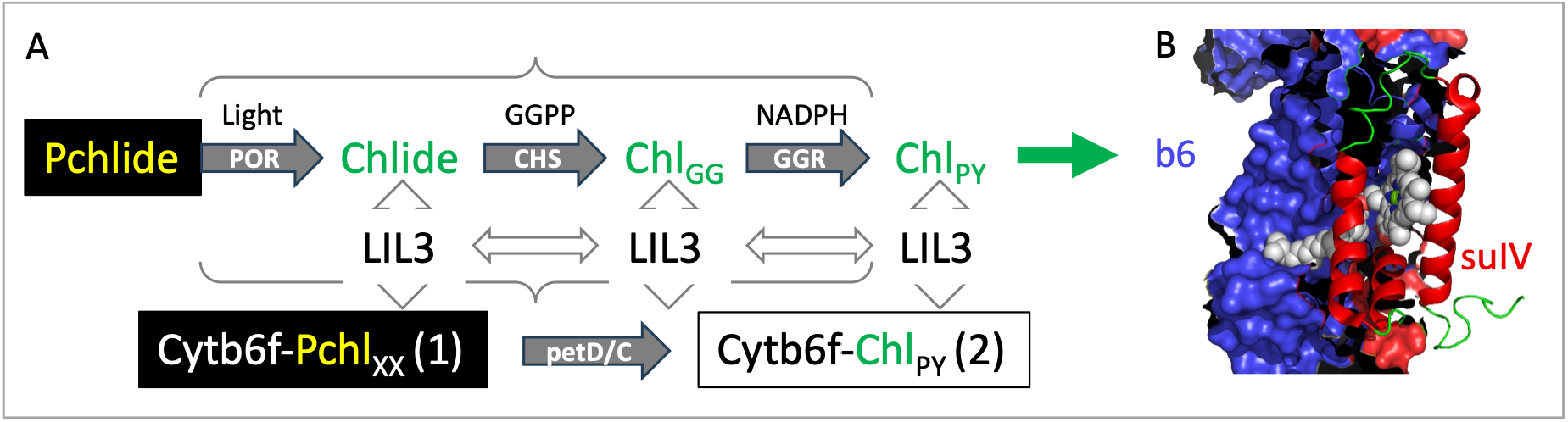
Model for regulation of Chl accumulation in Cyt b6f by LIL3. The enzymes protochlorophyllide oxidoreductase (POR, EC 1.3.1.33), and geranygeranyl reductase (GGR, EC1.3.1.83) are stromal enzymes. The light-harvesting like protein LIL3, and chlorophyll synthase (CHS, EC 2.5.1.62) are membrane integral proteins of the etioplast. All four proteins were determined to interact in barley etioplasts during synthesis of phytylated Chl (Tanaka et al. 2010; Mork-Jansson et al. 2015a; Hey et al. 2017). POR binds protochlorophyllide a (Pchlide) and regulates reduction of chlorophyllide a (Chlide) in a light and NADPH dependent reaction (grey arrow,(Garrone et al. 2015)). Chlide was synthesized upon illumination from Pchlide by POR (Light) or chemically supplied to etioplast membranes and shown to bind to LIL3 (Fig. 5, 6). LIL3 was found either in a non-assembled free form in the absence and in a complex with POR and CHS upon binding of Chlide (Mork-Jansson et al. 2015a). CHS has been shown to bind GGPP or PYPP in etioplasts isolated in darkness (Schmid et al. 2002). Chlide is esterified to GGPP or PYPP in etioplasts upon illumination or upon supplementation of Chlide (Eichacker et al. 1996). Both Chlide/Chl binding to LIL3 and binding of esterified Pchlxx/Chl_GG_ and Chl_PY_ binding to Cyt b6f dimers (2) was demonstrated. Results indicate that LIL3 regulates the synthesis of Chl_PY_ enabling Chl binding to the Cyt b6 subunit in the Cyt b6f monomer and Cyt b6f dimerization.

## Discussion

We show that LIL3-mediated conversion of Chlide to Chl and its subsequent release coordinate with the transition of Cyt b6f from monomer to dimer. Chl to Cyt b6f is prioritized during the first minutes of de-etiolation preceding detectable Chl incorporation into PSI/PSII cores (Fig. 2). A biogenesis of Chl-Cyt b6f is consistent with the requirement for rapid establishment of cyclic electron flow and ΔpH regulation before linear flow becomes dominant (Tikhonov 2014). The measured Kd of 246.6 ± 37 nM for Chlide-LIL3 further supports a high-affinity chaperone function analogous to cyanobacterial HliD (Proctor et al. 2020).

We find binding of Pchl to Cyt b6f and assembly of subunit IV (petD) in its dimeric complex in etioplasts. Etioplasts are not metabolically dormant and the identification of NDH, Cyt b6f, and of functional ATPase complexes indicate operational electron transfer in an etiorespiratory chain (Guéra et al. 2000). High rates of oxygen consumption have been measured in isolated etioplasts (Shevela et al. 2016), fully consistent with etiorespiration via the NDH–PQ–PTOX pathway during skoto-morphogenesis (Peltier et al. 2002). This etiorespiratory (chlororespiratory) electron flow, from stromal NAD(P)H via the NDH complex to the plastoquinone (PQ) pool and onward to PTOX, consumes O₂ while generating a modest proton-motive force (ΔpH/Δψ) across the prothylakoid membranes. This force can power, for example, Tat-dependent import of lumenal proteins into the prolamellar body and prothylakoids (Guéra et al. 2000; Mori and Cline 2001; Peltier et al. 2002; Shevela et al. 2016). Whether etiorespiration uses the Cyt b6f Q-cycle for generating ΔpH/Δψ is unclear; however, maintenance of the PQ pool in an oxidized state is essential for phytoene desaturase activity in carotenoid biosynthesis (Carol et al. 1999). For linear electron flow, Plastocyanin (PC) is present in etioplasts (Plesnicar and Bendall 1973), yet higher-plant plastids, unlike cyanobacteria, lack a terminal cytochrome oxidase downstream of PC (Peltier et al. 2010), creating a “PC dead-end” that precludes linear dark flow through Cyt b6f to O₂. We determined that the high rates of oxygen consumption in etioplasts were gradually compensated by an increasing oxygen production rate within the first four hours of deetiolation and assembly of the photosystem machinery (Shevela et al. 2019). Findings indicated the continuous operation of an electron transfer chain in etioplasts before and after the initiation of de-etiolation (Shevela et al. 2016). However, the function of Pchl-Cyt b6f dimer in etioplasts is unresolved (Guéra et al. 2000; Mori and Cline 2001; Peltier et al. 2002; Shevela et al. 2016).

### The Pchl binding to Cyt b6f

According to x-ray crystallographic structure (Stroebel et al. 2003), and biochemical analysis (Pierre et al. 1995; Mork-Jansson and Eichacker 2019), Cyt b6f binds one Chl a molecule per monomer, and Chl binding in Cyt b6f has been evolutionary conserved in oxygenic photosynthesis (Baniulis et al. 2013; Cardona 2019; Malone et al. 2019). Chl binding to Cyt b6f has been linked to the regulation of Cyt b6f subunit assembly, the regulation of electron flow by Cyt b6f, and the regulation of light-harvesting phosphorylation in state transition (Hasan et al. 2014; Tikhonov 2014). We showed that during de-etiolation, the binding of Chl was determined within seconds after the onset of Chl synthesis (Fig. 2, Cyt b6); while in etioplasts, Pchl was bound (Reisinger et al. 2008a). Quantitation of pigments in etiolated seedlings revealed a Pchl content of about 3-5 % relative to the dominant amount of Pchlide (Schoch et al. 1977). Hence, for the 6–7.5 × 10⁶ Pchlide molecules present, approximately 1.8–2.2 × 10⁵ Pchl molecules were present per etioplast (Eichacker et al. 1996). Interestingly, a similar amount of 1,5–2 x 10^5^ Cyt b6f complexes per etioplast was determined using differential spectroscopy (Mark-Aurel Schöttler and Lutz A. Eichacker, unpublished). Hence, Pchl and Cyt b6f appeared present in a ratio of about 1 in etioplasts. This suggests that all Pchl forms determined in the etioplast are bound to Cyt b6f, and the light induced spectroscopic changes in Cyt b6f dimers when corrected for the about three-fold lower specific extinction coefficient of Pchl relative to Chl, confirmed the 1:1 ratio for the Pchl to Chl exchange (Supplementary Figure 3). The finding that both esterified chlorins form part of the Cyt b6f structure during the transition from skoto-to photomorphogenesis, together with the quantitative correlation between Pchl/Chl and Cyt b₆f, suggests coregulation of Chl biosynthesis and the expression/assembly of Cyt b6f dimers. The finding that incompletely hydrogenated geranylgeraniol forms were bound to Cyt b6f further raised the question about the regulation of chlorin binding, and its functionality in the etioplast and how the hydrogenation affects an efficient exchange against Chl_PY_ when phytylated Chl becomes available to initiate the building of the photosynthetic electron transfer chain.

### High turnover of SUIV during Cyt b6f biogenesis in the etioplast

For functionality of the dimeric Cyt b6f complex, the chlorin binding site between subunits Cyt b6 and SUIV is of key importance. We find Cyt b6 is strongly binding to Pchl and Chl, and that the Cyt b6f core is not a static complex assembly unit. Radiolabeling of nascent membrane protein subunits during polysome run-off experiments in barley etioplasts show a high rate of petD expression, via incorporation of 35S-Met (Suppl. Fig. 2B). The corresponding SUIV protein accumulates in the dimer Cyt b6f complex during radiolabel chase kinetics. Interestingly, accumulation was found independent of the onset and the quantity of Chl synthesis (Suppl. Fig. 2A and B). The selectively high turnover of SUIV could indicate that this protein serves a housekeeping function for chlorin binding. In this scenario the stable Cyt b6 subunit forms the functional core of the complex, stabilizes SUIV, and thereby supports subsequent Cyt b6f assembly. In mature chloroplasts, SUIV likewise exhibits one of the highest turnover rates among all plastid proteins consistent with a dynamic role for the protein complex function beyond scaffolding (Li et al. 2023). Here, the function of SUIV is seen in expression regulation during Cyt b6f assembly. Translation of the Cyt f subunit has been described to be controlled by SUIV utilizing the control by the epistasy of synthesis mechanism (CES). In the CES mechanism, Cyt f that has assembled with the Cyt b6/SUIV core is withdrawn from its translation-initiation site on the petA mRNA, thereby allowing Cyt f expression (Choquet et al. 1998). In addition, the dedicated assembly factors NTA1/DEIP1 may be of key importance to understand the key subunit interactions for Cyt b6f core assembly (Sandoval-Ibáñez et al. 2022; Li et al. 2023).

### Cyt b6f maturation precedes photosystem biogenesis during de-etiolation

The function of Pchl binding to the Cyt b6 subunit in Cyt b6f besides scaffolding may be directly related to photosystem biogenesis. Pchl could act as a placeholder for Chl, ensuring rapid binding once Chl becomes available and thereby allowing the Cyt b6f complex to precede reaction-center biogenesis (Reisinger et al. 2008a). We found that within the first 10 s of illumination in-vivo or in-vitro, Chl accumulates exclusively in Cyt b6f and in LIL3 complexes with no Chl binding to PSI or PSII cores (Fig. 1). The binding of the newly synthesized chlorins to the LIL3 proteins appears catalytic (Fig. 2), while the Cyt b6f complex accumulates Chl during the reaction kinetics. The exclusive accumulation of Chl in Cyt b6f immediately after the onset of phototransformation could result from several molecular mechanisms that operate before the overall rate of Chl synthesis increases. As a consequence delivery of Chl_PY_ for assembly with the Chl binding photosystem complexes is delayed while during the start-up phase, functional assembly of Cyt b6f is prioritized to precede functional photosystem assembly. To date, no dedicated Chl chaperone for Cyt b6f has been found in cyanobacteria or mature chloroplasts, where the conserved HliD module was described to bind Chl, associate with ChlG and Ycf39, and deliver Chl to PSII (Chidgey et al. 2014; Proctor et al. 2020; Wysocka et al. 2025). In angiosperm etioplasts, we identified LIL3 to perform an analogous early-delivery Chl function, but show that it is the pre-assembled Cyt b6f dimer which is ready to bind the newly synthesized Chl. We also find that PSII shows Chl independent pre-assembly of D2 in three defined RC pre-complexes (Fig. 2A, RC_AI_). However, no Chl binding can be detected with assembly of pD1/D1 at the level of the RC complexes. In contrast, it required about 1 h in-vivo for Chl binding to be detectable in RC47 and RC monomer core subunits (Fig. 2B). The de-etiolation analysis therefore distinguishes the LIL3 function during early greening from HliD in Synechocystis, and BCM1/EGY1 for Chl homeostasis in mature leaves and from DEIP1/NTA1 regulating subunit assembly in chloroplasts (Wang et al. 2020; Li et al. 2023; Fu et al. 2025).

In our analysis, the etioplast LIL3 system appears to resemble an ancient pigment-chaperone module which was co-opted for rapid de-etiolation in land plants. The mechanism ensures a staged photosystem biogenesis to be initiated with Chl-binding to Cyt b6f. The ontogenetic presence of a functional Pchl-Cyt b6f dimer in darkness and its immediate transformation to a Chl-Cyt b6f dimer before any of the photosystem complexes bind Chl may mirror the phylogenetic evolution of the photosystem electron transport chain in general (Cardona 2019). We conclude that evolution of photosynthesis centers around the functional optimization of utilizing the chlorin molecules. We find an about 1/1 ratio of Pchl/Chl in the absence/presence of Chl, directly upon the onset of de-etiolation, indicating a Pchl exchange in the Cyt b6f complex (Suppl. Fig. 3). Here, the low selectivity of the Cyt b6/SUIV core for the reduction state of the chlorins isoprenoid side chain with the potential for both Pchl and Chl to be bound in their GG, PY, and intermediate forms exposes the question whether the reduction state of the isoprenoid chlorin side chain in Cyt b6f is part of the biogenetic regulation of the photosynthetic electron transfer chain. Here, the immediate establishment and maintenance of phytylated Chl binding during Cyt b6f assembly could be the key requirement for functional photosystem biogenesis.

### The Chl dependent Cyt b6f maturation and regulation of electron flow

We wondered whether the phytylated Chl-Cyt b6f precedes assembly and functional integration of light-harvesting complexes with the RC cores to control the onset of electron flow during de-etiolation or whether the immediate Chl-Cyt b6f assembly serves additional regulatory purpose (Shevela et al. 2016, 2019). In chloroplasts, the mobile redox cofactor plastoquinone (PQ) is of key importance for regulation of the Cyt b6f specific electron flow (Kambakam et al. 2016a). Binding of reduced plastoquinol (PQH2) to the Qp site is the decisive step for Cyt b6f specific electron divergence in linear electron transfer (LET) and the Q-cycle (Hasan et al. 2014). Both PQ molecules at the anterior and posterior Qp sites in the two Cyt b6 subunits of the Cyt b6f dimer are positioned adjacent to the phytyl chain of Chl a, indicating that the Chl a phytyl side chain may interfere with PQ and thereby with electron flow (Malone et al. 2019). A direct interaction between PQ and the chlorins isoprenoid side chain suggests that tuning the chemical structure and flexibility of the isoprenoid and ensuring its continuous integration into the Cyt b6f core could be of key importance for a differential regulation of electron flow in etiorespiration and photosynthesis. Here we find that, in addition to petD, the small plastid-encoded subunits petG/L/N also turn over in chloroplasts (data not shown). This indicates a requirement for continued protein expression and reassembly of petD/G/L/N to maintain Chl-specific Cyt b6f core assembly during photosynthesis.

This dynamic requirement for continuous Cyt b6f core assembly is consistent with the characterization of the *nta1/deip1* mutant in *Arabidopsis*. NTA1 (also designated DEIP1) encodes a conserved integral thylakoid membrane protein that functions as a dedicated assembly factor for the Cyt b6f complex. In the nta1 knockout mutant, loss of NTA1 (“new tiny albino 1”) causes a severe albino phenotype characterized by drastically reduced chlorophyll and carotenoid content, impaired thylakoid-membrane development, and seedling lethality under photoautotrophic conditions. BN-PAGE and immunoblot analyses revealed a specific and near-complete loss of the fully assembled Cyt b6f complex, while PSI and PSII core subunits (e.g. D1, CP43, PsaB) accumulate to levels comparable to wild-type during the first 6 h of illumination (ie. the early greening phase of dark-grown seedlings). Only thereafter do PSI/PSII levels decline, and light-harvesting chlorophyll-binding proteins (LHCPs) fail to accumulate substantially, resulting in the albino phenotype (Sandoval-Ibáñez et al. 2022; Li et al. 2023). These phenotypes are consistent with our kinetic evidence in barley etioplasts that Chl binding and maturation of the Cyt b6/PetD core precede functional integration of the photosystems during de-etiolation.

A subsequent suppressor screen then revealed an unexpected functional interplay between NTA1/DEIP1 and the cyclic electron flow protein PGR5 (Penzler et al. 2024). Double *pgr5 nta1/deip1* mutants are viable and accumulate detectable levels of the Cyt b6f complex, indicating that the strict requirement for NTA1/DEIP1 can be bypassed in the absence of PGR5. These findings suggest that NTA1/DEIP1 may primarily protect or stabilize the Cyt b6f complex against deleterious effects exerted by PGR5 (for example, excessive donor-side limitation or destabilization under fluctuating redox conditions), rather than acting as an absolutely essential assembly factor. The severe albino phenotype and the specific impairment of Cyt b6f accumulation in the single *nta1/deip1* mutant under standard conditions reinforces the importance of dedicated, dynamic regulation of the Cyt b6/PetD core and of the associated small subunit (PetG, PetL, PetN) assembly during the early phases of chloroplast biogenesis and de-etiolation. The de-etiolation-induced expression of NTA1/DEIP1 further underscores that this regulatory module is activated precisely when the etioplast-to-chloroplast transition requires prioritized Cyt b6f maturation.

The albino phenotype further indicates that the absence of a functional Cyt b6f complex in the *nta1/deip1* background directly limits light-induced activation of the full Chl-biosynthesis pathway. In *Chlamydomonas*, defects in the Cyt b6f complex (particularly at the Qₒ site) abolish light-induced expression of nuclear genes encoding tetrapyrrole/Chl-biosynthesis enzymes, demonstrating a retrograde signal coordinates Chl supply with electron-transport-chain assembly (Shao et al. 2006). In higher plants, similar retrograde control via the redox state of the plastoquinone pool, by Cyt b6f itself, and reactive oxygen species have been proposed to explain the secondary reduction of PSI/PSII and LHCPs in *nta1/deip1* (Li et al. 2023). In etioplasts, the modest proton-motive force (ΔpH/Δψ) generated by the NDH–PQ– PTOX etiorespiratory chain is already operational in darkness and might support Tat-dependent protein import and early thylakoid maturation (Guéra et al. 2000; Peltier et al. 2002; Shevela et al. 2016). PGR5-dependent cyclic electron flow around PSI, however, is light-activated and has not been shown to contribute to etiorespiration or dark ΔpH formation. The *pgr5 nta1/deip1* viability therefore raises the intriguing possibility that the primary role of NTA1/DEIP1 is to fine-tune Cyt b6f–PGR5 interactions that safeguards lumenal acidification and ATP synthesis once linear and cyclic electron transport commence.

Interestingly, PGR5 was also shown to participate in etiorespiration in etioplasts which provides additional mechanistic depth. In dark-grown Arabidopsis seedlings, PTOX mediates electron flow from NAD(P)H to O₂ via both the NDH complex and a PGR5-dependent pathway, thereby maintaining an oxidized PQ pool required for carotenoid biosynthesis and generating a modest ΔpH/Δψ across prothylakoid membranes (Kambakam et al. 2016b). Genetic evidence comes from the immutans (im) mutant (defective in PTOX), in which pgr5 suppresses the dark-accumulated phytoene phenotype and restores Pchlide levels in an additive manner with NDH defects. Thus, PGR5 is already active in etioplasts and may exert its deleterious effect on an incompletely assembled Cyt b6f complex already during skotomorphogenesis. The viability of the pgr5 nta1/deip1 double mutant (Penzler et al. 2024) could therefore reflect not only light-dependent protection of Cyt b6f but also an altered dark redox/ΔpH balance that relaxes the requirement for NTA1/DEIP1 during the etioplast-to-chloroplast transition. Future analyses of Pchlide accumulation, POR activity, and ΔpH formation in pgr5 and nta1/deip1 etioplasts will be critical to resolve whether the primary regulatory signal for staged Cyt b6f maturation and Chl biosynthesis is the Cyt b6/PetD core itself, its assembly factor NTA1/DEIP1, or downstream PGR5-dependent redox/ΔpH cues to regulate photosystem biogenesis in land plants.

### Model for a LIL3-orchestrated Chl hand-over to Cyt b6f

There is a single Chl *a* molecule that has to be bound per Cyt b6f monomer. Binding requires the protein pocket formed between subunits Cyt b6 and SUIV, so that the phytyl chain is positioned near the Qₚ site between the F and G helices of SUIV (Malone et al. 2019). We find a comobility of LIL3 and the Cyt b6f dimer in native PAGE, but have no direct LIL3 mediated Chl delivery to the photosystem complexes or to the Cyt b6f dimer. In contrast, we find LIL3 complexes binding both Chlide and Chl indicating that release of Chl from LIL3 and binding to photosystem proteins is regulated via the interaction between the isoprenoid side chains and the photosystem host proteins. As shown here for the Cyt b6/SUIV conformation this transfer principle could have evolved as it has to be permissive for chlorin turnover at multiple Chl binding sites. In the case of the Cyt b6f complex, the high rate of petD and of petG, L, and N expression, the strong binding of Chl to Cyt b6, and the Chl independent assembly of SUIV are indicating a strong biogenetic anchoring of this molecular mechanism.

Our model proposes that LIL3 increases the efficiency of Chl synthesis by keeping Chlide and Chl bound throughout the enzymatic modifications of the side chain. This could allow its release for direct Chl binding and most likely in exchange for membrane lipids to binding to the photosystem protein chains on demand (Solymosi and Mysliwa-Kurdziel 2021). In etiolated plants, POR is reducing Pchlide to Chlide in a light- and NADPH-dependent reaction. The binding of Chlide to LIL3 (Fig. 7) accelerates Pchlide binding to POR and hence phototransformation. However, the strongest increase in phototransformation efficiency is achieved in the presence of GGPP when esterification of Chlide is increased and a high quantity of the phytylated Chl can be bound to the Chl binding photosystem protein complexes. This indicates that LIL3 acts as chaperone binding Chlide and Chl. LIL3 positions the propionic acid side chain of Chlide relative to GGPP bound by membrane-integral CHS for efficient esterification. With the binding of GGR to the LIL3-CHS complex, the GGR mediated reduction to Chl-PY is effectively achieved while the chlorin remains bound to the LIL3–CHS–GGR supercomplex (Fig. 10). The stable association of Chlide and Chl with LIL3 appears as the basis for the fluorescent Chlorin-LIL3 complexes of increasing molecular weight (Mork-Jansson et al. 2015b). Mass-spectrometry analysis clearly indicates that additional protein subunits present in the fluorescent bands are related to photosystem assembly, as shown here for the Cyt b6f complex. We therefore postulate that Chlide/Chl remains bound to LIL3 throughout chlorin processing and that Chl is released from the complex in the presence of recipient binding pockets as shown for the prioritized receptor protein Cyt b6.

The LIL3-orchestrated Chl hand-over to Cyt b6f thus couples pigment biosynthesis to the etiorespiratory-to-photosynthetic switch in de-etiolation, ensuring that the central electron/proton relay is functional within seconds of illumination while the organelle remains protected by etiorespiration in darkness. In summary, our data support a regulatory pathway where LIL3 binds Chlide (Kd = 246.6 ± 37 nM), facilitating esterification and reduction, for delivery of Chl to Cyt b6f for Pchl exchange and dimerization. This process precedes photosystem biogenesis while the Pchl-Cyt b6f dimer sustains etiorespiratory proton pumping and PQ redox poising in darkness, highlighting LIL3’s dual role in biosynthesis/allocation and the key importance of early Cyt b6f assembly for both etiorespiration and the rapid onset of photosynthesis.

### Convergent reaction-center strategies in PSII and cytochrome b6f: Cpn60-dependent folding and continuous D1/SuIV expression

In addition to the prioritized accumulation of Chl in the Cyt b6f dimer, our 2D BN-PAGE/SDS-PAGE radiolabeling experiments (Fig. 2A) uncover a notable involvement of the plastid Cpn60 (GroEL/S) chaperonin during de-etiolation. Newly synthesized CP43 and CP47 show strong radiolabel accumulation at the position of the GroEL/S complex in the native dimension, with characteristic horizontal extensions and smearing in the second dimension indicative of transient chaperone–client interactions. This observation is independently supported by high-confidence CPN60 interactomics in *Chlamydomonas reinhardtii*, which identified both PsbB and PsbC as genuine CPN60/CPN20 substrates (Ries et al. 2023). Given that translation of these multi-spanning membrane proteins occurs on membrane-bound polysomes, the soluble stromal Cpn60 complex is envisioned to capture their large hydrophilic stromal domains at the membrane surface, assisting folding prior to Chl binding and integration into larger PSII subcomplexes.

This Cpn60 dependence underscores a key evolutionary divergence. While the reaction centers of PSI and PSII are structurally homologous and derive from a common ancestral homodimeric Type II reaction-center progenitor (Cardona 2019), the PSII lineage underwent genomic splitting of ancestral large core genes into the four distinct plastid genes *psbA* (D1), *psbD* (D2), *psbB* (CP47), and *psbC* (CP43). This fragmentation necessitated a strictly modular assembly pathway and additional chaperone layers, including Cpn60 for the stromal domains of CP43 and CP47, to accommodate the continuous high-turnover D1 repair cycle. Strikingly, this modular logic converges with that of the cyt b6f complex. In both systems a central Chl-binding “mold” subunit, D2 in PSII or Cyt b6 in Cyt b6f, is pre-synthesized and stabilized early in etioplasts, while a high-turnover “closer” subunit (D1 or SuIV/PetD) completes the functional reaction-center unit upon Chl availability (Plücken et al. 2002). Here, we find that the accessory subunits CP43/CP47 or Cyt f and small Pet subunits are incorporated subsequently.

The constitutive and rapid turnover of both D1 and SuIV, observed here in etioplasts independently of Chl synthesis and previously documented in mature chloroplasts, raises a fundamental mechanistic question: why must these two specific subunits be expressed continuously while the other subunits of the respective complexes remain stable? Protein synthesis is energetically costly, yet D1 and SuIV are among the most rapidly turned-over plastid-encoded proteins. The classical explanation for D1 turnover is ROS-mediated photodamage at the PSII reaction center; for SuIV, however, the equally high turnover rate— even in darkness—cannot be explained solely by oxidative damage. One possibility is that rapid subunit replacement serves not, or not only, to repair damaged protein but to enable exchange or renewal of the bound chlorophyll, and possibly of other cofactors, within the tightly integrated reaction-center Chl-binding pocket. The single Chl *a* per Cyt b6f monomer sits precisely at the Cyt b6/SuIV interface near the Qₚ site; similarly, D1 coordinates multiple Chl molecules and the non-heme iron in the PSII RC. However, D1/D2 precomplexes can form in etioplasts in the complete absence of Chl, and Chl is incorporated post-translationally into the RC (Müller and Eichacker 1999; Plücken et al. 2002; Knoppová 2022). Likewise we show that Chl and its precursor Pchl bind primarily to Cyt b6, and not to SuIV. This indicates that Chl exchange is regulated to occur independently from subunit turnover. A higher thermodynamic stability of the Chl-bound to Cyt b6 and D2 (Plücken et al. 2002; Hasan et al. 2014; Malone et al. 2019) likely favors selective turnover of the “closer” subunits SuIV and D1 to reset the pigment pocket without disassembling the entire core.

For PSII turnover, ribosome pausing during D1 translation has been well documented at multiple sites and linked to co-translational membrane integration, stromal-loop folding, and cofactor recruitment (Kim et al. 1991; Gawroński et al. 2018). In contrast, PetD (SuIV) insertion is post-translational and SRP/Alb3-dependent. Here, fully synthesized PetD is kept soluble by cpSRP54, delivered to the thylakoid, and inserted stepwise by Alb3 (Króliczewski et al. 2017). Hence Alb3, the chloroplast Oxa1/YidC homolog, functions in both co-translational (D1, Cyt b6) and post-translational (LHCII, PetD) modes, catalysing helix insertion rather than insertion of complete pre-folded proteins (Króliczewski et al. 2016). This hybrid strategy might allow precise stoichiometric control while avoiding stromal aggregation of hydrophobic subunits. Also, the continuous, organelle-localized synthesis of D1 and SuIV is consistent with the broader observation that certain high-turnover subunits of bioenergetic complexes remain encoded in organellar genomes. This pattern has been discussed in the context of the CoRR hypothesis (Allen 2017), although a detailed evaluation of organelle-genome retention lies outside the experimental scope of the present study.”

## Acknowledgements

We thank the Research Council of Norway (NFR335017) for financial support.

## Data/code availability

All data are contained in the manuscript and supplemental figures

