## Supplemental Figures for "Chlorophyll binding to Cytochrome b6f precedes photosystem I and II in barley etioplasts"

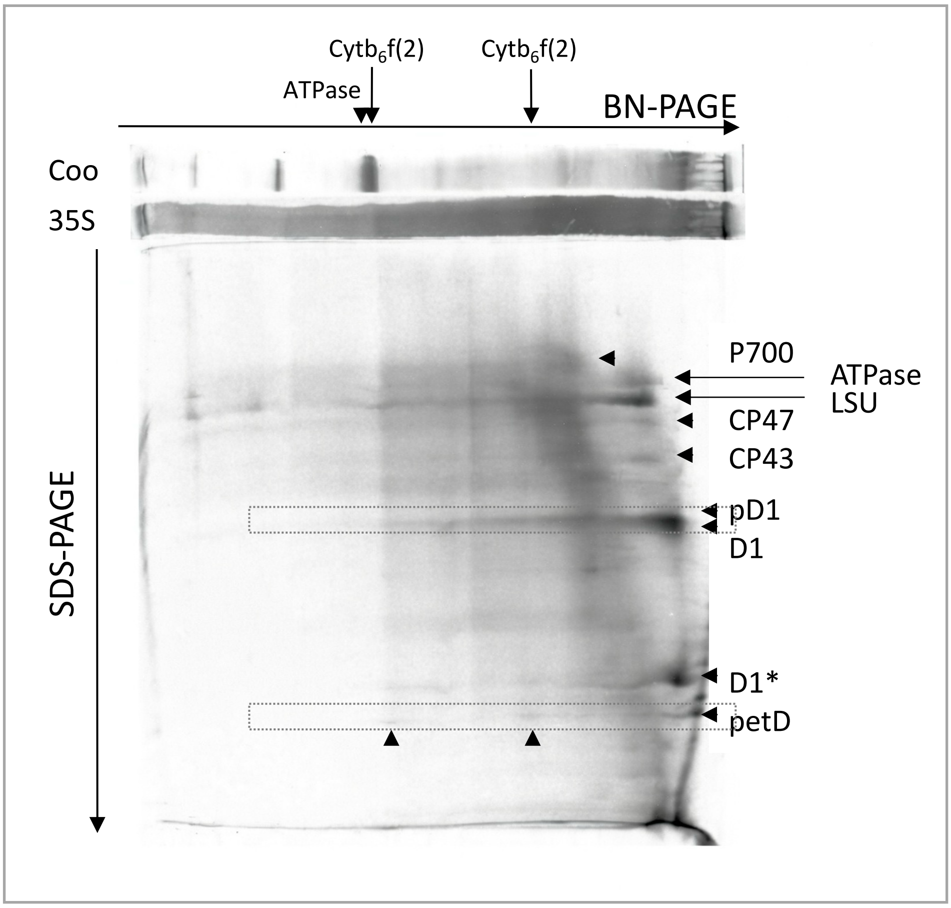


**Supplemental Figure 1: Visualization of etioplast membrane proteins by pulse radiolabeling**

Etioplast (1 x $10^8$) were isolated from 4.5 day-old dark-grown barley seedlings and were pulse-radiolabelled with 35S-Methionine in the dark for 15 min. Membranes were solubilized, and proteins separated by 2D Native-PAGE (BN-PAGE). Radiolabel in proteins was detected via Phosphoimaging. The location of radiolabelled protein subunits of PSI complexes (P700), and of PSII complexes (CP47, CP43, pD1, D1, D1*), of Cyt b6f complexes in monomer (1), and dimer (2) state (petD), of ATPase (a-, b-subunits) and of the large subunit of Rubisco (LSU) are marked in the 1D and 2D gel labelled. Protein subunits, and protein complexes were identified by gel blot and/or by MS analysis. The direction of protein mobility and protein bands in the 1D Blue native gel (BN-PAGE) are shown as horizontal gel slabs upon staining with Coomassie G250 (Coo) or radiolabel detection (35S).

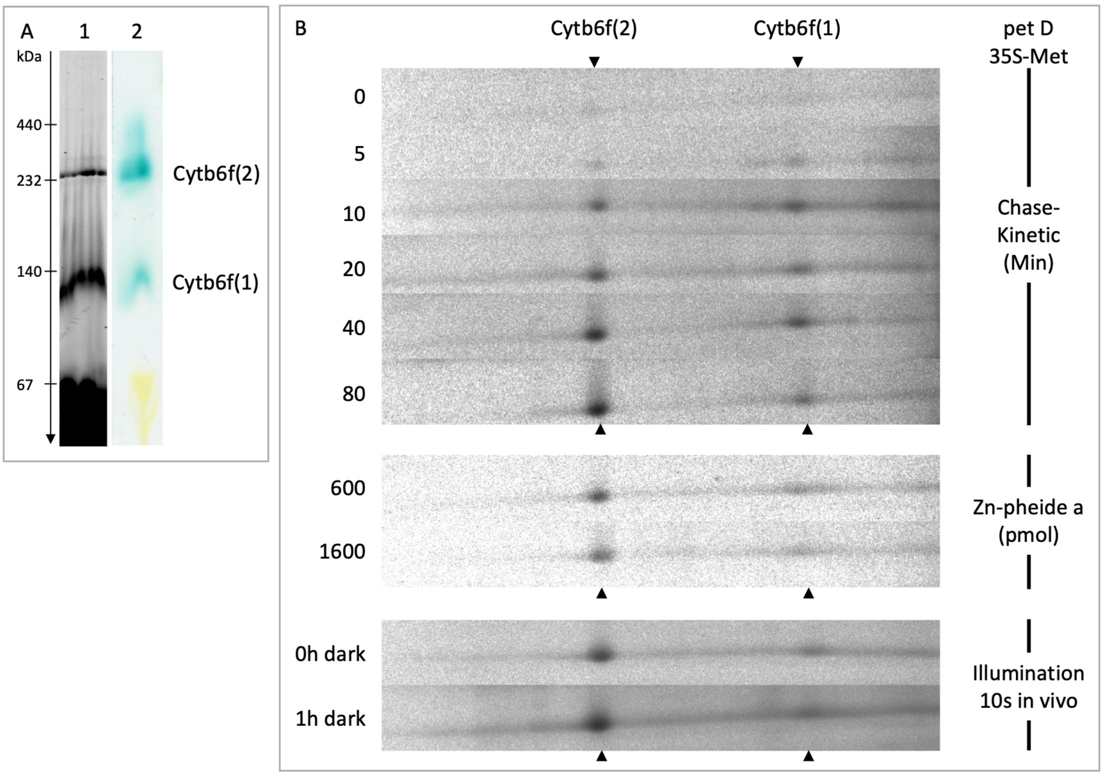


**Supplemental Figure 2: Determination of the Cyt b6f - complex in etioplast membranes**

Membranes from $10^8$ etioplasts were solubilized and protein complexes separated by CN-PAGE. Fluorescence of gels was recorded after laser excitation at 633 nm (A, lane 1). Peroxidase activity of cytochrome $b_6$ was probed by incubation of gels in TMBZ (A, lane 2). The two protein complexes show fluorescence at 670 nm and peroxidase activity. The complex also shows turnover of the petD gene product as shown by radiolabel incorporation (B, petD 35S-Met). The kinetic of radiolabel incorporation and maintenance of the petD expression and assembly in the two Cyt b6f complexes in the absence (Chase kinetic, min), and presence of Zn-phe synthesis (Zn-pheide a, pmol), and illumination of the etiolated plants in-vivo (illumination 10s in vivo) is shown (B). The petD protein product and other subunits of the Cyt b6f dimer were identified by mass spectrometry.

A

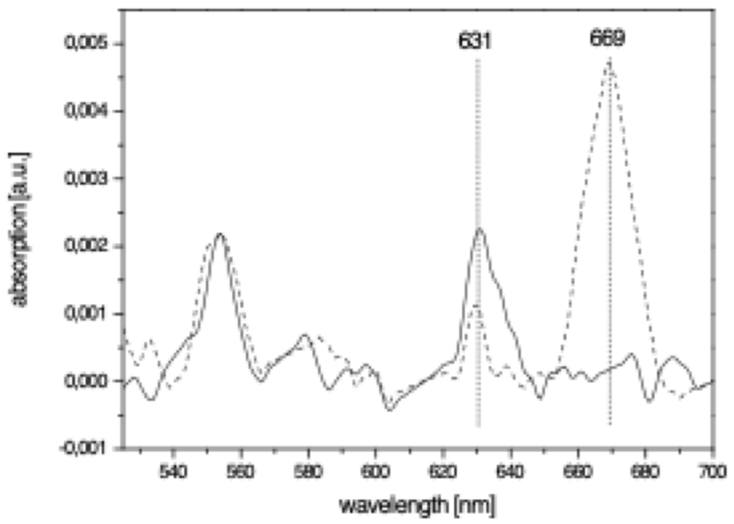


B

| Sample | Pchl (area) | Chl (area) | Chl/Pchl (ratio) |
|---|---|---|---|
| Dark | 0.03203 | - | |
| Light | 0.00849 | 0.07561 | |
| Total | 0.02354 | 0.07561 | 1.2 |

**Supplemental Figure 3: Determination of the Chl/Pchl ratio in the Cyt b6f dimer**

Replacement of Pchl with Chl in the dimeric Cyt b6f complex upon induction of Chl synthesis in etioplasts. Isolated etioplasts ($2 \times 10^8$) were incubated with 11 nmol GGPP for 15 min at 25 °C either in darkness (solid line) or after a 2-s white-light illumination (dashed line). Membranes were solubilized and protein complexes separated by LN-PAGE. The dimeric Cyt b6f complex ($F_{Y1}$ band) was excised and absorption spectra were recorded from 525–700 nm. Spectra were normalized to the absorption maximum of Cyt f at 554 nm (Bishop et al. 1972). The Cyt b6f complex isolated from dark-kept etioplasts showed an absorption maximum at 631 nm, corresponding to bound Pchl (Reisinger et al. 2008). Upon illumination in the presence of GGPP a maximum at 669 nm (Chl) appeared at the expense of the 631 nm peak. Traces of Pchl remained detectable, indicating that pigment exchange was not complete after 15 min of in organello Chl synthesis. Relative amounts of Pchl and Chl were determined from the spectral areas between 615–647 nm and 650–690 nm, respectively. Using the extinction coefficients of Pchl (30.4 $mM^{-1}$ $cm^{-1}$ in 80 % acetone at 626 nm; (Brouers and Michel-Wolwertz 1983) and Chl (82 $mM^{-1}$ $cm^{-1}$ in 80 % aqueous acetone at 663 nm; (Mackinney 1941), the area ratio [Chl]/[Pchl] ≈ 3.2 corresponded to a stoichiometry of approximately 1 : 1.2. This indicates that the single Pchl molecule bound per Cyt b6f monomer in etioplasts is replaced by Chl upon light-induced Chl synthesis.

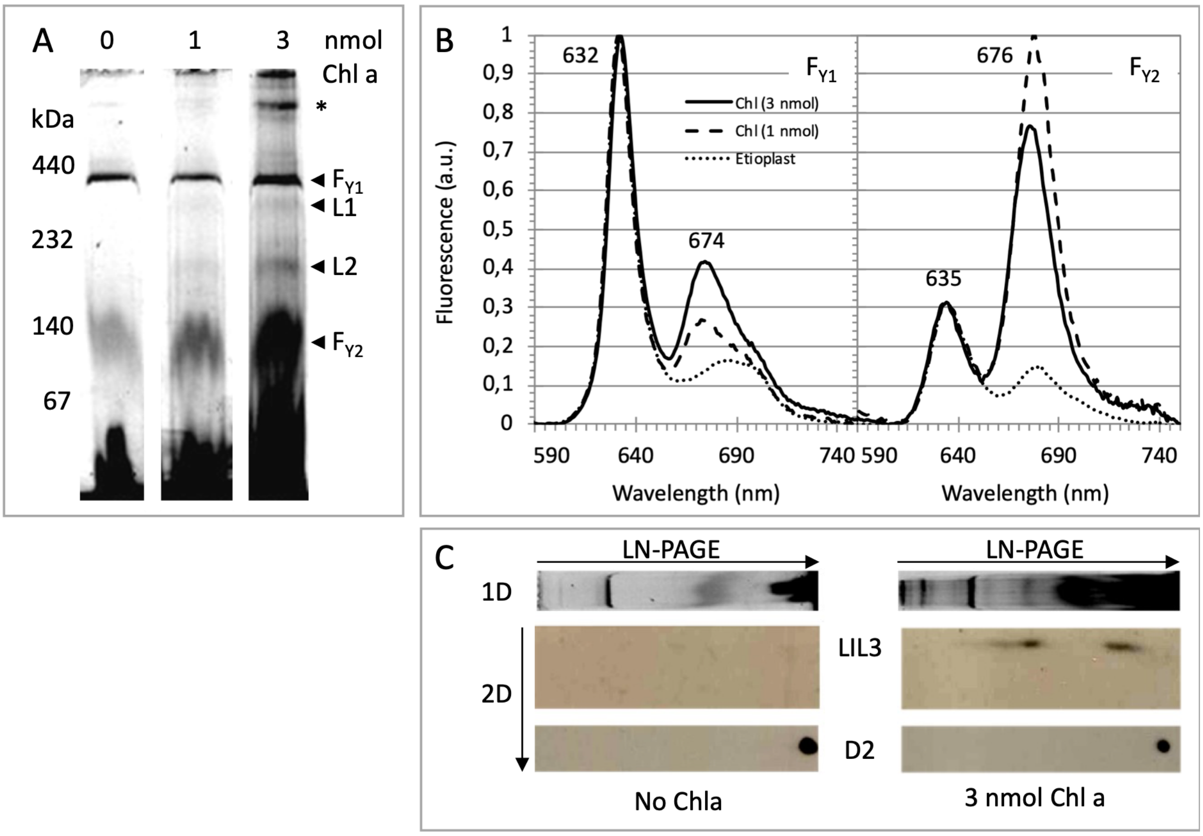


**Supplemental Figure 4: Accumulation of Chl in Cyt b6f dimer in etioplast membranes supplemented with Chl**

Plastids from dark-grown barley-seedlings ($10^8$) were isolated in green safe-light and incubated in the absence (A, lane 0) or presence of 1 or 3 nmol of Chl a, for 20 min (A, lane 1 and 3). Protein complexes were separated by native LN-PAGE, and fluorescence was recorded upon excitation/emission scanning at 633/670 nm (A). The location of fluorescent dimeric ($F_{Y1}$) and monomeric ($F_{Y2}$) Cyt b6f gel-bands and of fluorescent LIL3 gel-bands, $F_{L1}$, and $F_{L2}$, in the native gel are labelled top down. An unidentified protein complex is marked with a star (*). Fluorescent gel-bands at positions $F_{Y1}$ and $F_{Y2}$ were cut from the gel (A, lanes 0, 1, and 3 nmol Chl a) and the fluorescence emission spectra recorded from 590-750 nm using 440 nm for excitation. Spectra were normalized at 632 nm for gel-bands $F_{Y1}$ and at 635 nm for gel bands $F_{Y2}$ (B). The location of LIL3 and of D2 in the first-dimension native PAGE (LN-PAGE) was determined by gel-blot analysis of the LN-PAGE stripes after 2D-PAGE separation (2D) of the protein subunits for the two of the experimental settings (no Chl a, 3 nmol Chl a) utilizing antibodies against LIL3 and D2 (C).
